# Angicin, a novel bacteriocin of *Streptococcus anginosus*

**DOI:** 10.1101/2021.09.08.459082

**Authors:** Verena Vogel, Richard Bauer, Stefanie Mauerer, Nicole Schiffelholz, Christian Haupt, Gerd M. Seibold, Marcus Fändrich, Paul Walther, Barbara Spellerberg

## Abstract

As a conserved defense mechanism, many bacteria produce antimicrobial peptides, called bacteriocins, which give a colonization advantage in a multispecies environment. Here the first bacteriocin of *Streptococcus anginosus,* designated Angicin, is described. *S. anginosus* is commonly described as a commensal, however it also possesses a high pathogenic potential. Therefore, understanding factors contributing to its host colonization and persistence are important. A radial diffusion assay was used to identify *S. anginosus* BSU 1211 as a potent bacteriocin producer. By genetic mutagenesis the background of bacteriocin production and the bacteriocin gene itself were identified. Synthetic Angicin shows high activity against closely related streptococci, listeria and vancomycin resistant enterococci. It has a fast mechanism of action and causes a membrane disruption in target cells. Angicin, present in cell free supernatant, is insensitive to changes in temperature from −70 to 90 °C and pH values from 2-10, suggesting that it represents an interesting compound for potential applications in food preservation or clinical settings.

## Introduction

*Streptococcus anginosus* is commonly regarded as a commensal of mucosal membranes. It is found in the oral cavity, the gastrointestinal and the urogenital tracts^1,2^. However, increasing evidence points to the pathogenic potential of *S. anginosus.* The species can cause severe purulent infections at all body sites. Furthermore, isolation has been reported from invasive infections, abscesses, blood cultures and cystic fibrosis patients^3–7^.

In a natural environment, bacteria are part of multispecies communities, where space and nutrients are a limiting factor^8^. For efficient host colonization, it is important for bacteria to outcompete rival species. A mechanism for dealing with competing species is bacteriocin production. Bacteriocins are small antimicrobial peptides produced by bacteria to inhibit the growth of other, often closely related, bacteria^9,10^. These cationic peptides are ribosomally produced and either undergo major (class I) or nearly no post translational modification (class II). Class I and II bacteriocins are characterized by being thermostable and by possessing a molecular mass of less than 10kDa. The main mechanism of action is the disruption of cell wall and membrane integrity of target species. Bacteriocin producers are rendered insensitive towards their own bacteriocin by simultaneous expression of immunity proteins. Bacteriocin production is often regulated via quorum sensing (QS) mechanisms^11–14^. QS is a cell density dependent regulation of gene expression.

It is a common trait of lactic acid bacteria to produce bacteriocins, and several streptococcal bacteriocins have been described^15–17^. So far, bacteriocin production of *S. anginosus* has not been analyzed at great detail. Mendonca et al. investigated the regulation of a putative bacteriocin in *Streptococcus intermedius*^11^, which forms together with *Streptococcus anginosus* and *Streptococcus constellatus* the *Streptococcus Anginosus* Group (SAG). The *Streptococcal invasion locus* (*sil*), which is associated with the virulence of *Streptococcus pyogenes,* has also been shown to act as a regulator for bacteriocin production in *S. intermedius*^11,18,19^. The *sil* locus encodes a QS system consisting of a two–component system (SilA/B), a signaling peptide (SilCR) and an export/processing system (SilD/E). In *S. intermedius* the putative bacteriocin production was reported to be induced by the addition of SilCR, but the bacteriocin itself has not been addressed experimentally^11^. However, even though similar genes are present in *S. anginosus* neither bacteriocin production nor the *sil* locus has been investigated in this species.

To study the bacteriocin production of *S. anginosus*, we screened multiple *S. anginosus* strains for their ability to inhibit closely related bacterial species. Through targeted mutations, the genetic background for bacteriocin production of *S. anginosus* was identified. The novel bacteriocin was designated as Angicin and phenotypic traits like spectrum of activity, sensitivity towards heat, pH and enzymes as well as the mechanisms of inducing bacteriocin production are elucidated.

## Results

### Isolation of the bacteriocin producing *S. anginosus* strain BSU 1211

Producing bacteriocins is a common trait of streptococci. Screening 95 clinical *Streptococcus anginosus* isolates for strains inhibiting the growth of the closely related streptococcal species *Streptococcus anginosus* SK 52, *Streptococcus constellatus, Streptococcus intermedius* and *Streptococcus pyogenes* (data not shown) led to the identification of the strain *S. anginosus* BSU 1211. Out of the 95 clinical *S. anginosus* isolates 44 (46 %) were able to inhibit at least one of these streptococcal indicator strains, while strain BSU 1211 appears to be a potent bacteriocin producer that was active against multiple species and caused the biggest inhibition zone of all strains. When tested in a one-layer radial diffusion assay (RDA) it shows activity against other *S. anginosus* strains like the type strain SK52 and the species *S. constellatus, S. intermedius* and *S. pyogenes* (Fig. 1). In addition, the pathogen *Listeria monocytogenes* as well as other listerial species like *Listeria ivanovii* and *Listeria grayi* are strongly inhibited by *S. anginosus* BSU 1211 (Fig. 1). However, for both *L. monocytogenes* and *L. ivanovii* sometimes colonies could be seen in the inhibition zones. Subsequently, the clinically relevant ESKAPE pathogens, consisting of *Enterococcus faecium, Staphylococcus aureus, Klebsiella pneumoniae, Acinetobacter baumannii* and *Pseudomonas aeruginosa* were investigated for a susceptibility towards bacteriocins of *S. anginosus* BSU1211. These species cause nosocomial infections and are associated with high mortality^20,21^.

**Figure 1:**
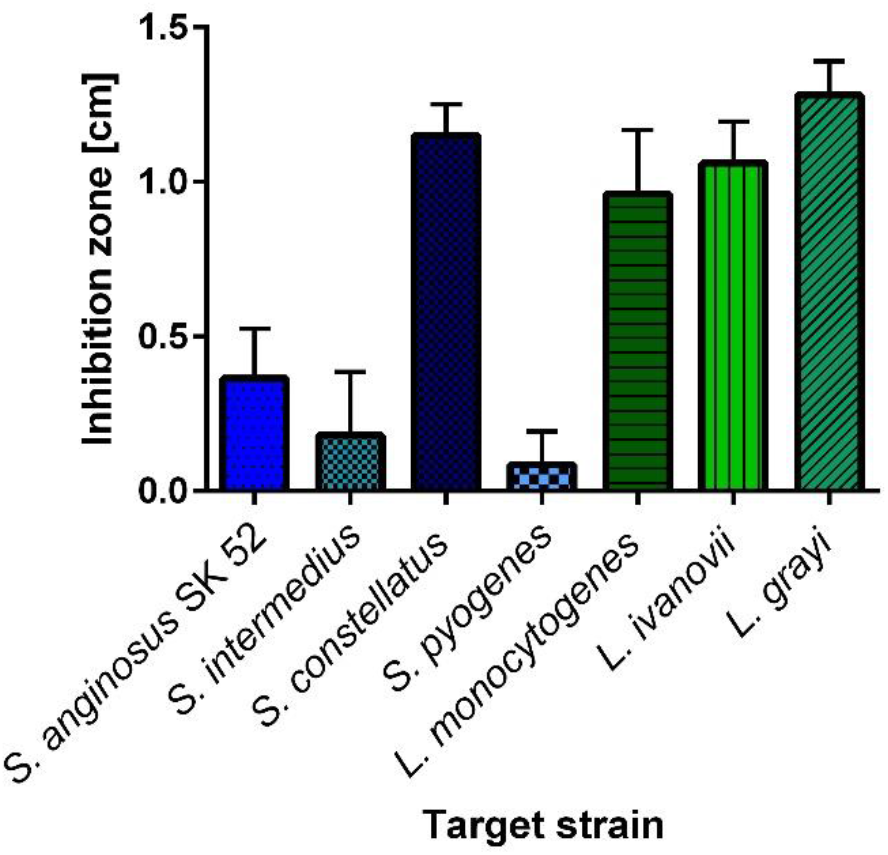
Spectrum of activity of *S. anginosus* BSU 1211. In a one-layer radial diffusion assay (RDA) *S. anginosus* BSU 1211 was tested for an effect against several target species. *S. anginosus* BSU 1211 shows activity against oral streptococci and several listeria species. Depicted is the mean ± standard deviation of at least five experiments.

However, no activity against any of these strains was seen. Additionally, neither staphylococci (*Staphylococcus aures, Staphylococcus epidermidis*), lactobacilli (*Lactobacillus gasseri, Lactobacillus acidophilus, Lactobacillus paracasei, Lactobacillus rhamnosus, Lactobacillus casei*), other streptococci (*Streptococcus agalactiae, Streptococcus mutans, Streptococcus dysagalactiae* subsp. *equisimilis, Streptococcus suis, Streptococcus porcinus, Streptococcus pneumoniae, Streptococcus mitis, Streptococcus oralis*), *Escherichia coli* nor *Bacillus subtilis* were inhibited by *S. anginosus* BSU 1211.

### Genetic background of bacteriocin production

To identify genes responsible for bacteriocin production in *S. anginosus* BSU 1211 the *sil* locus and the surrounding regions of the genome were sequenced. In *S. intermedius* and other species of the SAG, a putative bacteriocin encoding region is adjacent to the quorum sensing locus *sil*^11^. We found that all *sil* genes are present in *S. anginosus* BSU 1211 and that an adjacent region, resembling the putative bacteriocin accessory region described in Mendonca et al.^11^ (Fig. 2). The open reading frames (ORF) were designated bacteriocin like peptide3 (*blp*3) in accordance with similar findings *in S. pyogenes*^16^. Bioinformatic analysis of the genes in this region with BLAST (https://blast.ncbi.nlm.nih.gov/Blast.cgi) and Bagel4 (http://bagel4.molgenrug.nl/) predicted three putative class II bacteriocins (*blp3.1, blp3.4, blp3.6*) (Table 1)^22,23^. By Bagel4 core peptide analysis, the proteins encoded by *blp3.1* (MKTKTLEKFEVLNSEMLARVEGGGCNWGDFAKSGIAGGAGNRLRLGIKTITWQGVVAGAVGGAIIGKVGYGATCWW) and *blp3.6* (MNTKSLEKFEVLNSEMLASVEGGKIGAGEAAQALAVCTVAGGTIGSVFPGVGTAVGAILGAQYCTGAWAIIRAH) resemble the known bacteriocin BlpU (Table 1). Both contain GxxxG motifs, which are a hallmark for two peptide bacteriocins^24^. A high homology to the Bovicin variant 255 and Garvicin Q is found for Angicin (MKESENFMLNLKGEVVNELSDVQLSEISGGGSGYCKPVMVGANGYACRYSNGRWDYKVTKGIFQATTDVIVKGWAEYGPWIPRH) (encoded by *blp3.4*).

**Figure 2:**
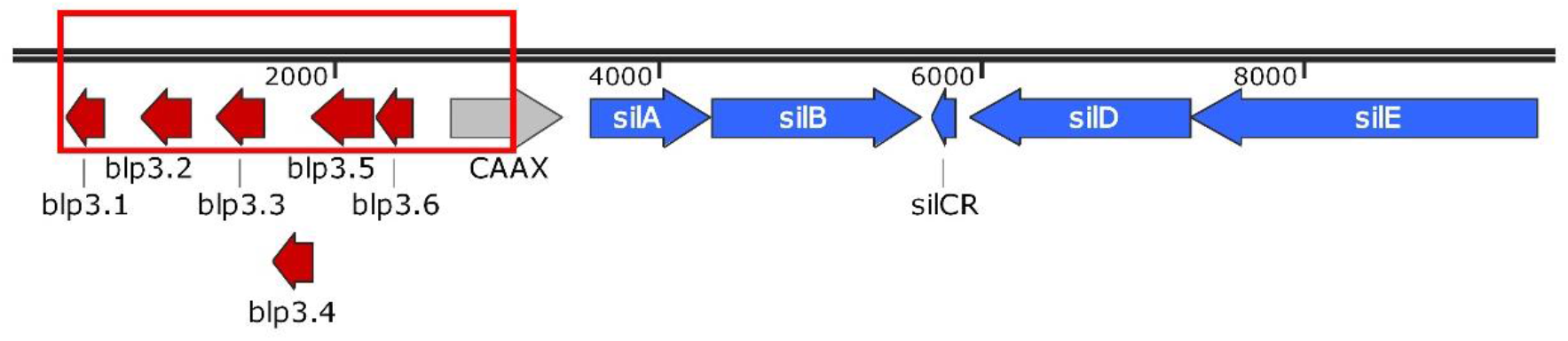
Genetic organization of *S. anginosus* BSU 1211. The *sil* genes and the *blp3* region were sequenced and open reading frames were adapted from Mendonca et al. and labeling was done in line with Hertzog et al. ^11,16^. In red the deleted region of *S. anginosus* BSU 1211Δ*blp3* is demonstrated. Schematic overview was constructed with SnapGene.

**Table 1:** Bioinformatic analysis of *blp3* region

|  | Bagle4 | BlastX (Query /Identiity) |
| --- | --- | --- |
| Blp3.1 | BlpU | bacteriocin [*S. anginosus*] (98 %/100 %) |
| Blp3.2 | No function determined | Hypothetical protein [*S. anginosus*] (99 %/99 %) |
| Blp3.3 | Putative piscicolin 126 immunity protein, Pisl | Bacteriocin immunity protein [*S. anginosus*] (98 %/100 %) |
| Blp3.4 | Bovicin variant 255 | MULTISPECIES: Garvicin Q family class II bacteriocin [*Streptococcus*] (90 %/100 %) |
| Blp3.5 | Bacteriocin self-immunity, putative | hypothetical protein SII_1169 [*S. intermedius* C270] (98 %/100 %) |
| Blp3.6 | BlpU | MULTISPECIES: bacteriocin [*Streptococcus*] (98 %/100 %) |
| CAAX | No function determined | MULTISPECIES: CPBP family intramembrane metalloprotease [*Streptococcus*] (99 %/99.95 %) |

The finding that *blp3.1, blp3.4* and *blp3.6* all encode a leader peptide with a double glycine motif, further supports the theory of these peptides being bacteriocins. All other deduced proteins did not match any of the known bacteriocin. Further sequence analysis with Bagel4 predicted *blp3.3* and *blp3.5* to encode immunity proteins. For *blp3.2* no hypothetical function could be determined, while *CAAX* showed high similarities (99 %) to CPBP family intramembrane metalloprotease in BLAST analysis.

The *blp3-*region could be identified in some completely sequence *S. anginosus* isolates (J4211, C1051, MAS624, C238, FDAARGOS_1155, FDAARGOS_1357) as well as in several other clinical *S. anginosus* isolates (BSU 1326, BSU 1370, BSU 1401). The *blp3*-regions of *S.* anginosus J4211, C1051, BSU 1370 and BSU 1401 contain the same genes and are 99 % identical with strain BSU 1211. Blp3.1 of *S. anginosus* J4211, C1051, BSU 1326 and BSU 1401 has one mutation (R42G) when compared to Blp3.1 of *S. anginosus* BSU 1211, while the deduced peptide sequence of *blp3.4* and *blp3.6* is identical in all analyzed strains.

The putative regulatory *sil* locus of *S. anginosus* BSU 1211 appears to be conserved in *S. anginosus* strains. The genes *silA, silD* and *silE* are identical to the respective genes described in the whole genomes sequence of *S. anginosus*_1_2_62CV. Furthermore, *silB* shows 99.85 % and *silCR* 99.32 % sequence identity. However, a point mutation was found in *silCR* leading to an amino acid change (F11L) in the mature autoinducing peptide SilCR, when compared to SilCR_SAG-B_. Thus, SilCR of *S. anginosus* BSU 1211 (GWLEDLFSPYFKKYKLGKLGQPDLG) is labeled SilCR_SAG-C_.

### Regulation and induction of bacteriocin production

Even though the strains BSU 1326, 1370 and 1401 encode the same bacteriocins as *S. anginosus* BSU 1211 no or only moderate inhibition of target strains can be observed in a one-layer RDA (Fig. 3). To investigate if bacteriocin production in these strains is dependent on the *sil* locus, synthetic autoinducing peptide SilCR was added in a one-layer RDA. Inhibition zones increase as soon as SilCR_SAG-C_ is administered to the bacterial culture (Fig. 3). Without the addition of SilCR_SAG-C_ *S. anginosus* BSU 1326 and 1370 are not able to inhibit any target strain. For *S. anginosus* BSU 1326 inhibition zones with a diameter of 0.36 cm for *S. constellatus* and 0.47 cm for *L. monocytogenes* were observed in the presence of 100 ng SilCR_SAG-C_ Similar effects occurred for the strain BSU 1370 and BSU 1401 (Fig. 3). However, the addition of SilCR_SAG-C_ did not lead to larger inhibition zones for *S. anginosus* BSU1211, suggesting that maximal bacteriocin expression is already present in this strain. Furthermore, it is possible that in strain BSU 1211 bacteriocins are expressed constitutively. The autoinducing peptide SilCR_SAG-C_ alone did not cause any inhibition of target strains.

**Figure 3:**
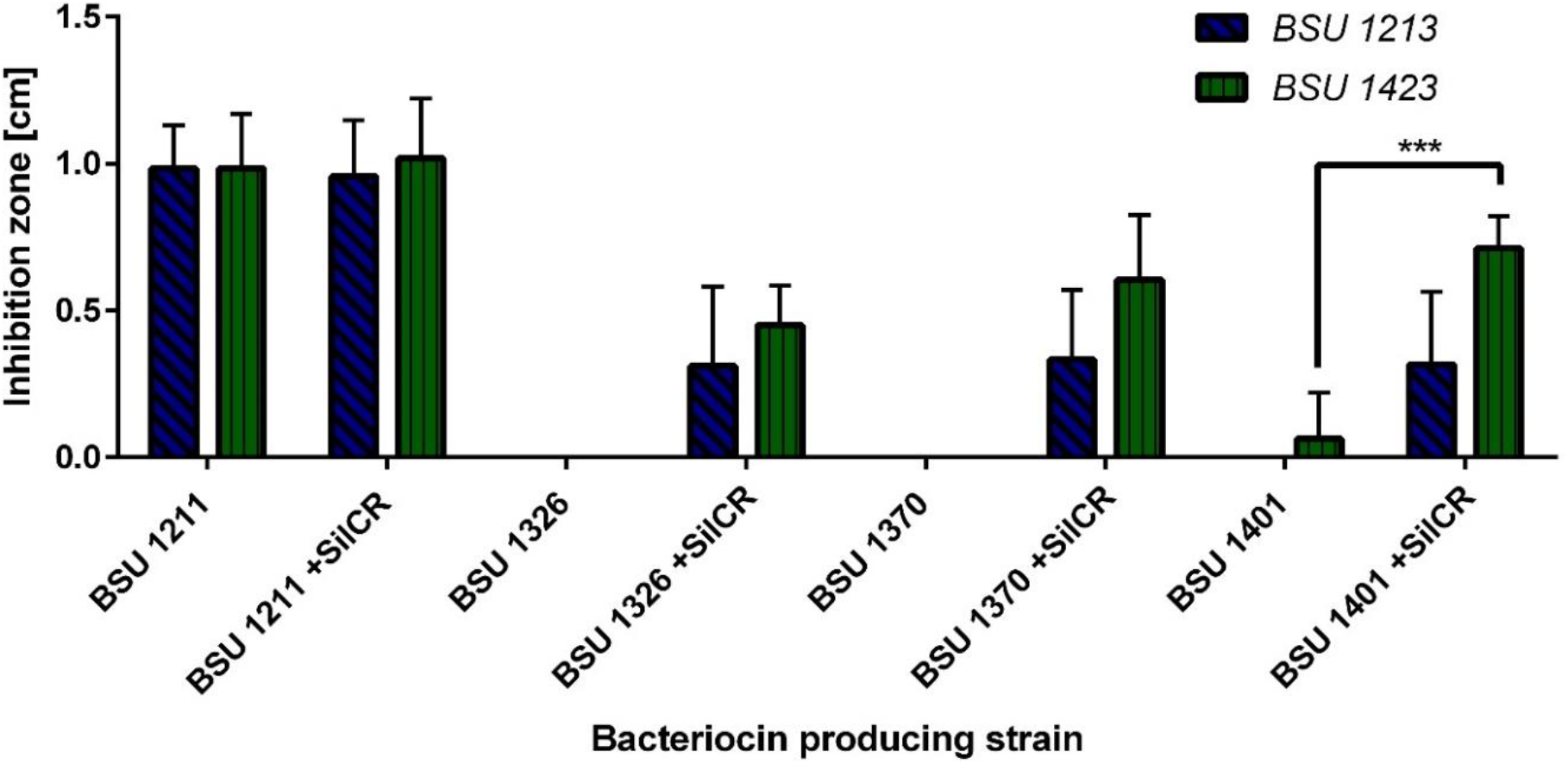
Induction of angicin production by addition of SilCR_SAG-C_. Result of a one-layer radial diffusion assay. *S. constellatus* BSU 1213 and *L. monocytogenes* BSU 1423 were used as target strains. The inhibitory activity of *S. anginosus* BSU 1211, 1326, 1370 and 1401 was determined either in the presence or absence of 100 ng SilCR_SAG-C_. Depicted is mean ± standard deviation of at least five independent experiments. A significant difference between SilCR_SAG-C_ supplemented and untreated *S. anginosus* strains was tested with a Mann-Whitney-U-test (*** indicates p<0.001).

### The *blp3* region

Often all genes necessary for bacteriocin production are clustered in one genetic region. To verify that in *S. anginosus* BSU 1211 the *blp3*-region is indeed responsible for the bacteriocin production and thereby the inhibition of target strains, a deletion mutant comprising the entire *blp3* region was constructed (*S. anginosus* BSU 1211Δ*blp3*) (Fig. 2). *S. anginosus* BSU 1211Δ*blp3* failed to inhibit any strain that was sensitive towards the bacteriocins produced by *S. anginosus* BSU 1211. The inhibition of these strains could be reestablished by complementing *S. anginosus* BSU 1211Δ*blp3* with a plasmid (pAT18-*blp3*) that encoded the deleted *blp3* region. Maximum activity of the complementation mutant against *L. monocytogenes, L. grayi* or *L. ivanovii* could be observed with adding synthetic SilCR_SAG-C_ (Fig. 4). As controls, the inhibition of *L. monocytogenes, L. grayi, L. ivanovii and S. constellatus* by *S. anginosus* SK52 (a strain which lacks the *sil* and *blp3* region) transformed with pAT18-*blp3* and *S. anginosus* BSU 1211Δ*blp3* transformed with the empty plasmid was investigated.

**Figure 4:**
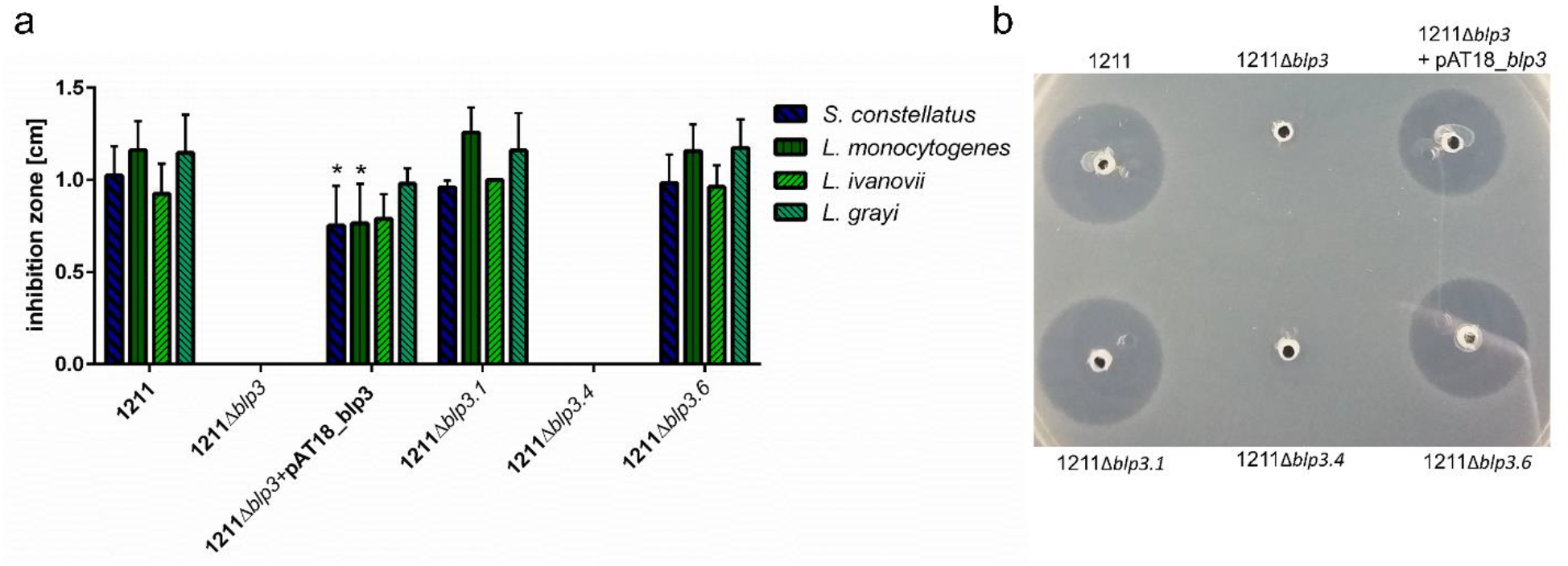
Inhibition of target strains by *blp3*-region mutants. (a)The effect of completely deleting *blp3* or solemnly deleting the predicted Angicin gens was tested in a one-layer radial diffusion assay against the four most prominent target strains *S. constellatus, L. monocytogenes, L. grayi, and L. ivanovii.* Furthermore, a *S. anginosus* BSU 1211Δ*blp3* was complemented with the *blp3*-region encoded on plasmid pAT18. The complementation mutant was supplemented with 100 ng SilCR_SAG-C_. (b) Result of a one-layer RDA against *L. grayi.* Shown is the mean ± standard deviation of at least five independent experiments. Significant differences between mutant and wildtype strain were calculated with a Mann-Whitney-U-test (* indicates p<0.05).

Both strains never showed an inhibition of target strain. This data clearly shows that the *sil* as well as the *blp3* region are necessary for bacteriocin production, however the question arises which of the predicted bacteriocins is functional and responsible for the activity or if more than one Angicin is produced and active, like it is the case in other streptococci ^25,26^ or if the observed activity arises from synergistic effects of encoded peptides as described for Garvicin KS of *Lactococcus garvieae*^27^. To investigate this, single Angicin genes (*blp3.1, blp3.4* and *blp3.6*) were chromosomally deleted and subsequently these mutant strains were assessed for their ability to inhibit the four most prominent target strains (*L. monocytogenes, L. ivanovii, L. grayi, S. constellatus*). Unfortunately, BSU 1211Δ*blp3.1* contained an additional point mutation in a non-coding area. It is not suspected that this mutation has any influence on bacteriocin production. A deletion of *blp3.1* and *blp3.6* did not alter antimicrobial activity. Deleting *blp3.4* caused a complete loss of inhibition activity. Thus, suggesting that Blp3.4, subsequently labeled Angicin, is the bacteriocin responsible for the antimicrobial activity of *S. anginosus* BSU 1211.

### Angicin activity

Angicin is synthesized as an 84 amino acid long prepetide. Typically, class II bacteriocins possess a leader peptide, which is cleaved of during export, resulting in the mature and active peptide. The first 30 amino acids of the 84 amino acid long prepeptide form a double glycine leader peptide which after processing leads to a 54 aa long mature Angicin. The peptide sequence of Angicin was compared to similar bacteriocins, like Bovicin 255, Garvicin Q and BacSJ. While Bovicin variant 255 has a sequence identity of approximately 64 %, Garvicin Q and BacSJ only have approximately 42 % sequence identity (supplementary Fig. S1). To verify that Angicin is accountable for the antimicrobial activity of *S. anginosus* BSU 1211 it was synthetically produced. Angicin was synthesized without its respective leader peptide and without any post translational modifications. The experimental mass of synthetic Angicin was 6053.1 Da, as determined by mass spectrometry, closely matching the theoretic mass of this 54 amino acid peptide (6052.9 Da).

The isoelectric point of the peptide was determined at 10.2 by PSL Heidelberg (https://www.peptid.de/), while computational analysis of the amino acid sequence with Bachem gave an isoelectric point at 9.6 and a net charge of 4 at neutral pH (https://www.bachem.com/service-support/peptide-calculator/). The synthetic peptide was dissolved in ultra-pure water, and its activity was examined in a two-layer RDA against *L. monocytogenes*. Concentrations of 50 μg/ml Angicin caused an inhibition zone of 1.09 ± 0.16 cm, and even concentrations of 1.56 μg/ml Angicin were still able to inhibit growth of *L. monocytogenes* (Fig. 5a). We then tested as to whether or not Angicin shows the same spectrum of activity as the strain *S. anginosus* BSU 1211. Based on a two-layer RDA (Fig. 5b) we find that strains that *L. monocytogenes, L. grayi*, *L. ivanovii* and *S. constellatus,* which are susceptible towards *S. anginosus* BSU1211 (Fig. 1), are also sensitive towards Angicin. However, the inhibition zones of *S. constellatus* were not completely clear in all experiments. *S. anginosus* SK52 showed a similar inhibition zone size like *S. constellatus*. However, the inhibition zones of *S. constellatus* were not completely clear in all experiments. *S. anginosus* SK52 showed a similar inhibition zone size like *S. constellatus*. In a second step, we investigated the Angicin susceptibility of the clinically relevant and the ESKAPE pathogens.

**Figure 5:**
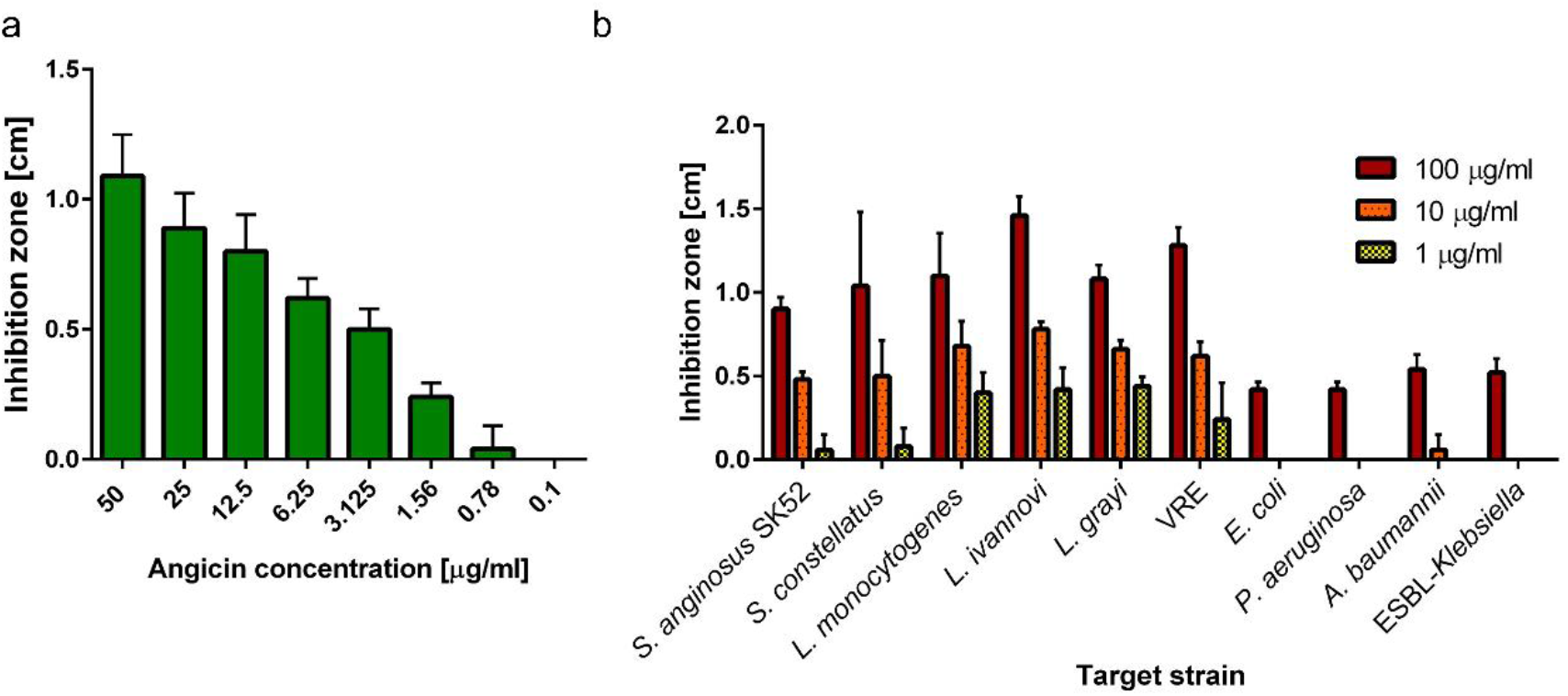
Effect of synthezied angicin on target strains. (a) Different Angicin concentrations were tested for activity against *L. monocytogenes* in a two-layer RDA. (b) Angicin concentrations of 100, 10 and 1 μg/ml were tested against different bacterial species in a two-layer RDA. 6.05 μg/ml equal 1 μM Angicin. VRE indicates Vancomycin resistant *Enterococcus faecium*, ESBL is an abbreviation for extended spectrum beta-lactamase. Shown is the mean ± standard deviation of five independent experiments.

We find that some of these pathogens, such as *S. aureus, B. subtilis,* are completely insensitive towards Angicin, while others showed a moderate inhibition at Angicin concentrations of 100 μg/ml (*E. coli, K. pneumoniae, P. aeruginosa, A. baumannii*). Interestingly, amongst these Angicin-sensitive ESKAPE pathogens was vancomycin resistant *Enterococcus faecium* (VRE), which causes nosocomial infections and pose a huge health concern due to rising numbers and limited treatment options^28^. The tested VRE strain was as sensitive in our assay as the listeria species.

### Activity of cell free supernatant

Many bacteriocins are secreted into the culture supernatant. To better understand the physical and chemical properties of Angicin, cell free supernatant (CFS) of BSU 1211 was prepared. A two-layer RDA was carried out to assess the activity of CFS in comparison to the bacteriocin activity of bacterial cells from BSU 1211.

In these experiments the previously identified target strain *L. monocytogenes* showed sensitivity towards CFS (Supplementary Fig. S2). However, no clear inhibition zone was observed. To show that Angicin is indeed the active component in CFS of BSU 1211, CFS of *S. anginosus* BSU 1211Δ*blp3.4* was prepared and tested for activity. It showed no antimicrobial activity against *L. monocytogenes.* In a next step, the activity of CFS of BSU 1211 after exposure to heat or pH-changes was investigated. Results are summarized in Table 2 and demonstrate that the *S. anginosus* CSF can tolerate up to 1 h of 60 °C, an acidic pH of 2, and can be kept at −20 °C and −70 °C for one year without loss of activity. However, exposure to 100 °C for 30 min as well as treatment with proteinase K destroyed the bacteriocin activity of CFS. A treatment with lipase had no effect on CFS activity. As a control, CFS of *S. anginosus* BSU 1211Δ*blp3* was challenged with the same treatments, but no inhibition of target strains was observed with BSU 1211Δ*blp3* CFS. By contrast, we find that synthectic Angicin resembled the activity pattern of BSU 1211 CFS when exposed to the above-mentioned heat challenges and enzyme treatments. In contrast to BSU 1211 CFS, however, we find that synthetic Angicin caused clear inhibition zones of *L. monocytogenes* and it showed no reduced activity upon heating at 90 °C for 1h.

**Table 2:** Bacteriocin stability in cell free supernatant against *L. monocytogenes* after various treatments. “+” indicates no altered activity, “reduced” indicates reduced activity and “−” means a complete loss of activity, N.D. indicates not determined.

| Treatment |  | Activity CFS | Activity Angicin |
| --- | --- | --- | --- |
| **Temperature** | 60 °C- 1 h | + | + |
|  | 80 °C – 1 h | Reduced | + |
|  | 90 °C -1 h | Reduced | + |
|  | 100 °C – 30 min | - | N.D. |
| **pH** | 2 | + | N.D. |
|  | 4 | + | N.D. |
|  | 6 | + | N.D. |
|  | 8 | + | N.D. |
|  | 10 | + | N.D. |
|  | 12 | - | N.D. |
| **Proteinase K** | 1mg/ml | - | - |
| **Lipase** | 1 mg/ml | + | + |
| **Stability after 1 year** | -20 °C | + | N.D. |
|  | -70 °C | + | N.D. |

To investigate, if the Angicin activity of CFS can also be demonstrated in liquid culture experiments, we measured the growth of *L. monocytogenes* in the presence of 25 % CFS (supplementary Fig. S3). We find that *L. monocytogenes* is significantly inhibited in the presence of 25 % CFS from the wildtype strain BSU 1211(p-value: 0.0006-6h), while 25 % CFS from the deletion mutant *S. anginosus* BSU 1211Δ*blp3* is even slightly enhanced, when compared to growth in medium without CFS. Furthermore, the activity of Angicin was evaluated in a bacterial survival assay. Since 25 % CFS was the best concentration in an inhibition assay, the same concentration was chosen for a survival assay. In the survival assay not only *L. monocytogenes* but furthermore *L. grayi* and *L. ivanovii* were investigated. In phosphate buffer supplemented with the CFS of *S. anginosus* BSU 1211Δ*blp3* no reduced survival could be seen (Fig. 6). In contrast, incubation with CFS of *S. anginosus* BSU 1211 significantly reduced the number of surviving cells (Fig. 6). The effect on *L. monocytogenes* was a 10-fold reduction, the inhibition of *L. ivanovii* and *L. grayi* with a 10^4^-fold and 10^6^-fold reduction respectively was even more pronounced. In a next step, listerial cells treated with 25 % CFS of *S. anginosus* BSU 1211 were analyzed via transmission electron microscopy to check for loss of membrane integrity (Fig. 6). For *L. monocytogenes* more damaged cells can be seen in the treated group than in the control group. It appears like numbers of *L. ivanovii* cells decrease after treatment with active CFS for 15 min. Furthermore, a lot of cell debris can be seen.

**Figure 6:**
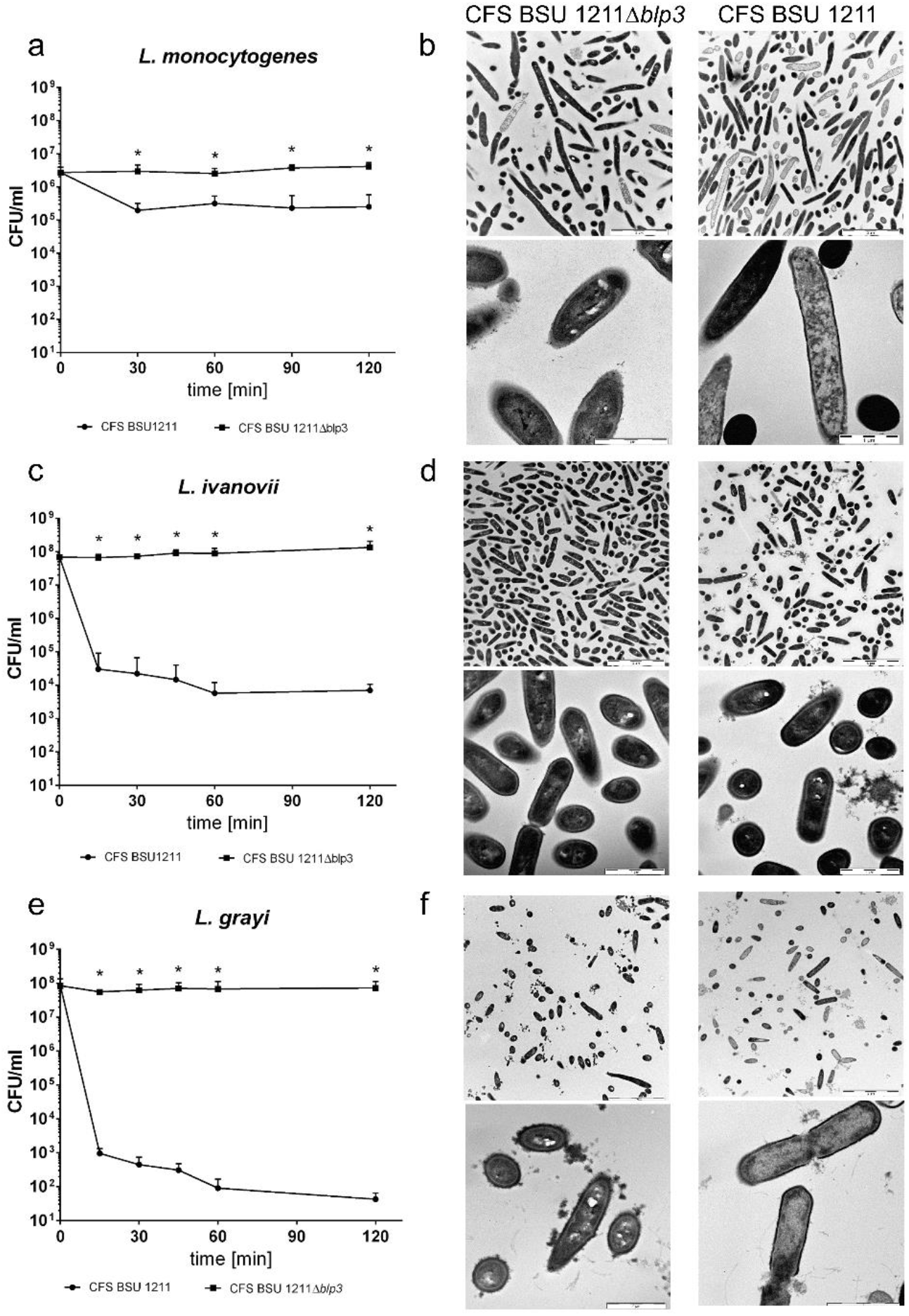
Effect of cell-free supernatant (CFS) of *S. anginosus* BSU 1211 or *S. anginosus* BSU 1211Δ*blp3* on *Listeria monocytogenes, L. ivannoi* and *L. grayi*. Survival of *L. monocytogenes* (a), *L. ivannovi* (c), *L. grayi* (e) in the presence of 25% CFS of *S. anginosus* BSU 1211 or *S. anginosus* BSU 1211Δ*blp3* over a time course of 2 h. Transmission electron microscopy (TEM) pictures of *L. monocytogenes* cells (b) treated with 25% CFS of *S. anginosus* BSU 1211 or *S. anginosus* BSU 1211Δ*blp3* for 2 h at 37 °C. TEM pictures of *L. ivannovi* (d) and *L. grayi* (f) were done after 15 min incubation with either 25% CFS of *S. anginosus* BSU 1211 or *S. anginosus* BSU 1211Δ*blp3*. Depicted for the survival assay is the mean + standard deviation of five independent experiments. TEM experiments were conducted once. Scale bar of the upper images is 5 μm, scale bar of the lower images represents 1 μm. Significant difference in survival were calculated with a Mann-Whitney-U-test (* indicates p<0.05).

Especially for *L. grayi* membrane disruption is visible in samples treated with active CFS for 15 min. Multiple cells appear lysed and loss of membrane integrity often seems to appear at sites of division. None of this can be seen in control cells treated with CFS of *S. anginosus* BSU 1211Δ*blp3*.

### Mechanism of action

The most common mode of action of bacteriocins is membrane permeabilization. To see if also Angicin uses this mechanism, a SYTOX Green Membrane Permeabilization Assay was conducted. SYTOX Green is a fluorescent DNA stain that can enter the bacterial cell after membrane disruption. Therefore, the fluorescence intensity is an indicator of membrane integrity. Treating *L. monocytogenes* with 100, 50 or 25 μg/ml synthetic Angicin lead to a significant SYTOX enrichment, indicating membrane disruption of the target cells (Fig. 7). Already after 5 min of incubation with either 100 or 50 μg/ml synthetic Angicin the membrane is measurably disrupted, indicating a very fast mechanism of action of the peptide (p-value: 0.0079). Additionally, no significant difference between the ethanol treated cells (positive control) and cells treated with 100 μg/ml for 15 min can be detected.

**Figure 7:**
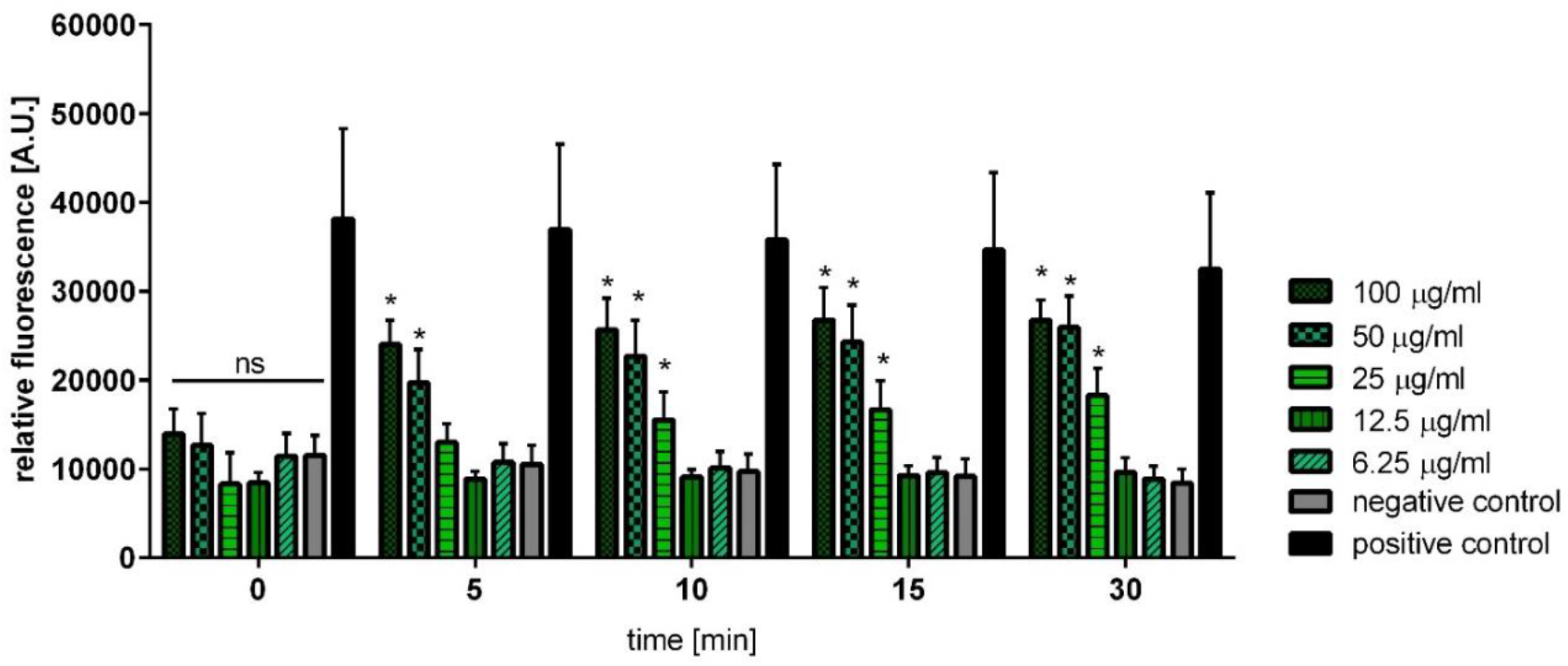
Effect of angicin on membrane integrity of *L. monocytogenes*. Bacterial cells were incubated with different Angicin concentrations for 1h at 37 °C and fluorescence was detected using Tecan Plate Reader (Excitation: 488 nm; Emission: 530 nm) at the indicated time points. 6.05 μg/ml Angicin equal a concentration of 1 μM Angicin. *L. monocytogenes* treated with 70 % Ethanol was used as positive control. Depicted are five independent experiments conducted with three technical replicates. Statistically significant differences to the negative control were calculated with Mann-Whitney-U test (* indicates p<0.05).

### Immunity proteins

For self-protection bacteriocin producers express immunity proteins. To explore, if Blp3.3 protein represents as predicted an immunity protein a functional assay was performed. The gene *blp3.3* was introduced into plasmid pAT28 under the control of a endogenous promotor or without a promotor and transformed into the Angicin sensitive target strains *S. anginosus* SK52 and *S. constellatus* BSU 1213. In a next step a one-layer RDA was performed and the effect of *S. anginosus* BUS 1211 on the target strains was determined. Inhibition zones of mutants were compared to inhibition zones of the wildtype strain harboring empty pAT28 plasmid. Transformation of *S. anginosus* SK52 with *blp3.3* under the control of an endogenous promotor led to a significantly reduced sensitivity towards Angicin (Fig. 8). Whereas *blp3.3* without promoter did not alter inhibition zone sizes. Comparable results were obtained for the *S. constellatus* strain BSU 1213. Introducing the *blp3.3* gene with an endogenous promotor led to significantly decreased inhibition zones. However, introducing *blp3.3* alone into *S. constellatus* BSU 1213 had no significant effect. In summary, this data supports the function of Blp3.3 as an immunity protein.

**Figure 8:**
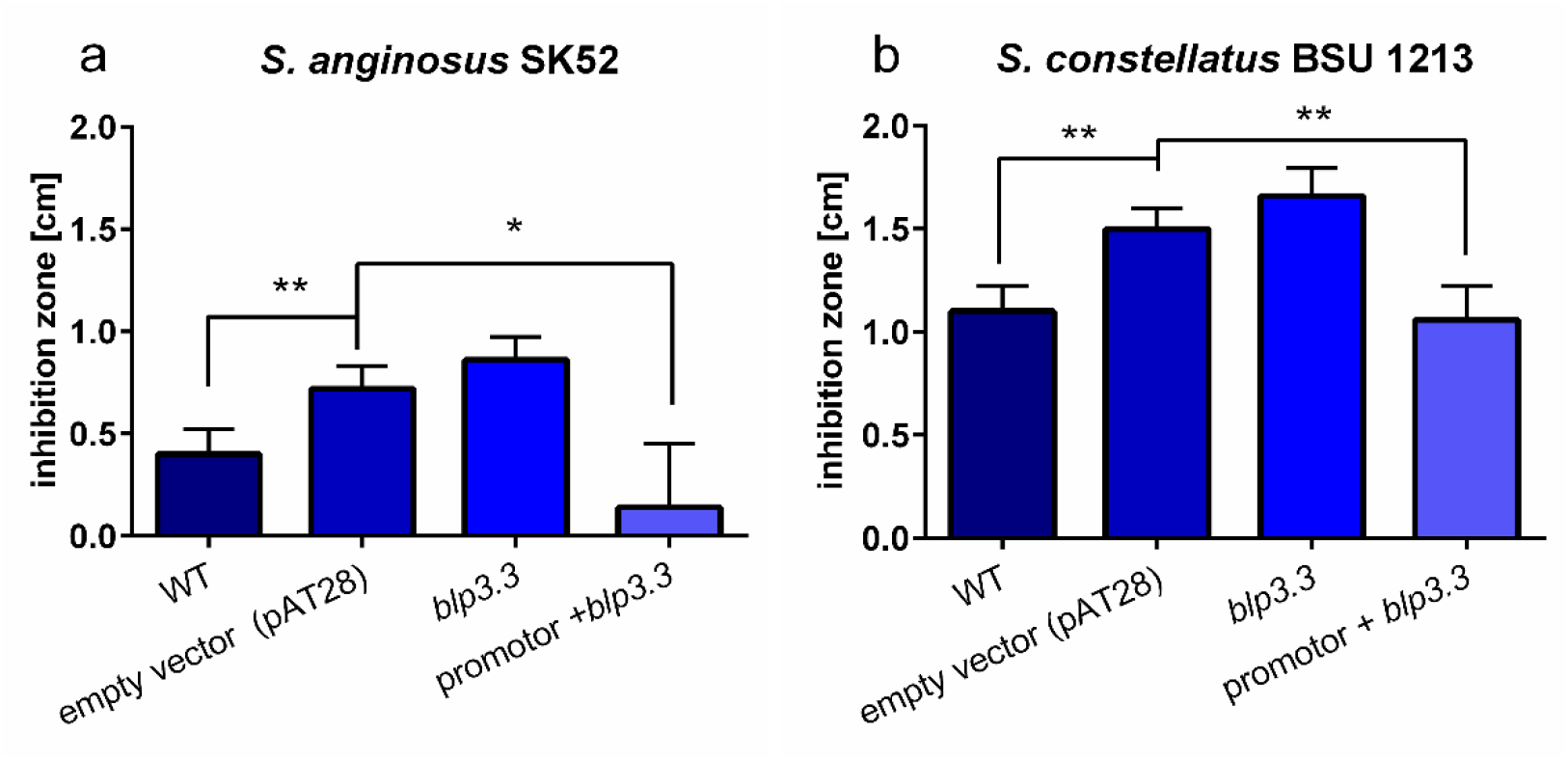
Influence of *blp3.3* on inhibition zone diameters. *Blp3.3* of S. anginosus BSU 1211 was transformed in susceptible target strains (a-S. anginosus SK 52, b-S. constellatus) either alone or under the control of an endogenous promotor. These mutants were used as target strains in a one-layer radial diffusion assay and tested against *S. anginosus* BSU 1211. Data was collected in at least five independent experiments. A significant difference to target strains transformed with the empty vector was tested with a Mann-Whitney-U-test (* indicates p<0.05, ** indicates p<0.01 and *** indicates p<0.001).

Subsequently, Blp3.3 protein was recombinantly expressed in *E. co*li and purified to investigate its effect on the antimicrobial activity of Angicin. To that end, we preincubated Blp3.3 together with Angicin for 1 h at 37 °C before the Angicin activity was tested in a two-layer RDA. We found that Blp3.3 alone already caused a zone of inhibited growth (supplementary Fig. S4) and it did not decrease Angicin activity on the target strain *L. monocytogenes*, irrespective of whether we used equimolar, substoichometric or superstoichometric concentrations of Blp3.3. Moreover, recombinant Blp3.3 protein did not affect the inhibitory activity of *S. anginosus* BSU1211 against *L. monocytogenes* when analyzing the inhibition zone size in a one-layer RDA. In addition, there was no inhibition of *L. monocytogenes* by Blp3.3 protein with or without *S. anginosus* BSU 1211Δ*blp3* (supplementary Fig. S4).

## Discussion

In their natural habitat, bacteria are part of complex multi-species environments. *S. anginosus* typically resides in the oral cavity which can be colonized with up to 700 different species^29,30^. Closely related species often need similar nutrients or inhabit a similar space, therefore there is an ongoing competition between these species. Secreting antimicrobial substances, like bacteriocins, provides the producing organism with a great colonization advantage^31,32^.

Putative bacteriocin production of streptococcal species from the *S. anginosus* group has previously been reported^33,34^. To elucidate the molecular background of this phenomenon, 95 *S. anginosus* strains were evaluated for inhibitory activity in coculture experiments, which led to the selection of strain BSU 1211 for its potent antimicrobial activity. In this strain the genetic basis of bacteriocin production and regulation was further investigated. Bacteriocin production is often regulated through quorum sensing systems^35^. In streptococci, like *S. intermedius* and *S. pyogenes* the *sil* system controls bacteriocin production^11,16^. A *sil* locus was also identified and predicted to be functional in many of the surveyed *S. anginosus* strains. In nearly all these strains inducing the *sil* two component system (SilAB) by addition of the autoinducing peptide SilCR_SAG-C_, turns on bacteriocin production. In *S. anginosus* the *sil* locus is directly adjacent to putative bacteriocin genes, encoded in the *blp3* region. By genetic mutagenesis we could show that this region is indeed responsible for bacteriocin production, since a complete deletion mutant of *blp3* was unable to inhibit the growth of any target strains. To identify the bacteriocin the three predicted bacteriocin genes of this locus were mutated separately. A deletion of *blp3.1* and *blp3.6* did not alter bacteriocin activity, while deleting *blp3.4* completely abolished the inhibitory effect. Interestingly, the deduced peptides of blp3.1 and blp3.6 both contain a GxxxG motif, which is a key feature of two peptide bacteriocins24, that function only as a couple. Typically, the structural genes of two peptide bacteriocins are located directly next to each other. But this is not the case in the *blp3* region of *S. anginosus*. It may be possible that by natural mutagenesis of this region the genes were separated from each other and are therefore no longer active. The putative bacteriocin region of SAG is a known hotspot of genetic diversity^11^. Possibly, the peptides are active in vitro when environmental conditions or other signals are present.

The identified *S. anginosus* bacteriocin was designated Angicin. The Angicin prepetide is 84 amino acids long of which the first 30 amino acids form the leader peptide. Mature Angicin has a predicted pI of 10.2 and a molecular mass of 6053 Da. In our experiments Angicin showed a good antimicrobial activity against *S. anginosus, S. constellatus, L. monocytogenes, L. grayi, L. ivanovii* and VRE. Moderate inhibitory activity could be observed against *E. coli, K. pneumoniae, P. aeruginosa, A. baumannii*.

Comparing peptide sequences, Angicin shows high homology to Bovicin (variant 255) and is grouped into the Garvicin Q family which also includes BacSJ as a member^36–39^. Garvicin Q is a class IId bacteriocin of *Lactococcus garvieae* with a wide spectrum of activity. It is active against all tested strains of carnobacteria, enterococci, lactococci, leuconostoc and listeria and most of the tested *Lactobacillus and Pediococcus* strains. While all species of the genera *Bacillus, Campylobacter, Staphylococcus* and *Streptococcus* as well as *P. aeruginosa, Salmonella typhimurium* and *Candida albicans* are not susceptible^40^ BacSJ is produced by *Lactobacillus paracasei* and shows a very similar spectrum of activity with the only exception being a resistance of *Lactococcus garvieae* against BacSJ but not Garvicin Q^38^. Bovicin (variant 255), a bacteriocin firstly isolated from *Streptococcus gallolyticus,* shows only a narrow spectrum of activity with inhibitiory properties against *E. faecium, Butyrivibrio fibrisolvens, Lactobacillus ruminis* and *Peptostreptococcus anaerobius.* All Gram negative species as well as *Streptococcus bovis, Bifidobacterium thermophilum* or *Ruminobacter amylophilus* are insensitive towards the peptide^39,41^. In summary, Garvicin Q and BacSJ seem to be mainly important for intra- and interspecies competition. This may also be true for Angicin, which inhibits besides other *S. anginosus* isolates also other streptococci, VRE and listeria.

*S. anginosus* BSU 1211 is not able to inhibit Gram negative species and Angicin only shows an inhibition of Gram negatives when administered at high concentrations. As already demonstrated for antibiotics, a reason for bacteriocin ineffectiveness against Gram negative species seems to be the outer membrane^42,43^. It inhibits an interaction between bacteriocin and its cognate receptor. Nisin binding to lipid II for example is inhibited by the outer membrane. However, inhibition of Gram negative species can be enhanced by the addition of chelating agents like ethylenediaminetetraacetic acid, that destroys the outer membrane^44,45^. For Angicin a similar situation may be the case.

Bacteriocins often kill target cells by membrane permeabilization. The electrostatic interaction between the cationic peptide and the negatively charged bacterial membrane is important as well as amphiphilic structures^46^. By a SYTOX Green membrane permeabilization assay it was demonstrated that Angicin leads to a disruption of the membrane in target cells. Already 5 min after Angicin treatment (Fig. 7) a significant difference in fluorescence signal to untreated cells is visible, indicating a fast mechanism of action. A rapid target cell lysis is not unusual and has also been observed in bacteriocins from enterococci and lactobacilli^47,48^. This data is supported by survival assays and TEM analysis, showing reduced cell numbers and rupture of membranes in listeria cells (Fig. 6). However, membrane permeabilization is mainly seen when high concentrations of Angicin are administered. For other bacteriocins it has been speculated that bacteriocins only work as membrane perturbers at concentrations that exceed what is present in a natural environment^49^. In lower or more natural concentrations, bacteriocin producers can use different mechanisms to inhibit competitors. Subtilisin for example interferes with quorum sensing of target organisms and thereby inhibits biofilm formation^50^. Thus, apart from membrane disruption other mechanisms may also play a role in the antibacterial activity of Angicin. For example, Garvicin A has been shown interfere with cell division by inhibition of septum formation ^51^. In CFS treated *L. grayi* cells loss of membrane integrity often appears at sites of division, which points in a similar direction.

Bacteriocin producers are rendered insensitive towards their own bacteriocin by simultaneous expression of immunity proteins^52^. Immunity proteins have a high specificity towards the bacteriocin and several different modes of action are already described, including loss of bacteriocin binding, sequestering, bacteriocin export or degradation of the bacteriocin^53–56^. By bioinformatic analysis of the *blp3* region, *blp3.3* was predicted as an immunity protein. It shows similarities to the immunity protein PisI, which protects against the class IIa bacteriocin piscolin 126^57,58^. Heterologous expression of *blp3.3* in Angicin target strains led to a decreased sensitivity of these strains. Similar results were obtained for PisI as well as for other bacteriocins^55,59,60^. However, there seems to be no direct interaction between blp3.3 and Angicin, because preincubation of both proteins with each other did not alter the activity of Angicin. Instead of a direct interaction immunity proteins may form a tertiary complex with the bacteriocin and the bacteriocin receptor^59,61,62^. Formation of this complex seems to depend on conformational changes after the initial binding of the bacteriocin to its receptor.

So far bacteriocins have been used in food preservation, as anticancer agents, treatment of peptic ulcer, as spermicidal agents, for skincare, in veterinary medicine and in agriculture^63,64^. Angicin inhibits *L. monocytogenes,* a species posing a major health risk for food safety^65^. For an application in food preservation, characteristics like stability over a wide pH and temperature range are important^66^. With remaining active in a pH ranging from 2-10 and at temperatures till 90 °C both criteria are met by Angicin. Furthermore, Angicin shows activity against VRE, a pathogen posing a great health risk^28^. This pathogen can cause infections like endocarditis, bacteremia and urinary tract infections and is one of the main causes of nosocomial infections^67,68^. Treatment options are limited and new ways to deal with this pathogen are needed^69^.

This is the first study describing and characterizing a bacteriocin of *S. anginosus.* The small catatonically charged Angicin not only inhibits closely related species, but furthermore the important food pathogen *L. monocytogenes* and clinically relevant vancomycin resistant *E. faecium.* This activity complemented with high thermal and wide pH stability make Angicin an interesting compound for food preservation and antimicrobial therapies.

## Methods

### Bacterial strains and growth conditions

All strains used in this study are summarized in supplementary Table S1. *Escherichia coli* was cultivated in lysogeny broth (LB-Miller). *E. coli* EC101 and DH5α were hosts for plasmids pAT28 and pAT18. *E. coli* mutants were incubated in the presence of 100 μg ml^−^ 1 spectinomycin (pAT28) or 400 μg mg ^−1^ erythromycin (pAT18). Liquid cultures were kept aerobically at 37 °C on a shaker (180 rpm), whereas plates were incubated at 37 °C and 5 % CO_2_. All other used bacterial strains were cultivated on sheep blood agar plates, for liquid broth THY medium (Todd-Hewitt Broth supplemented with 0.5 % yeast extract) was used. If necessary, streptococci were grown with 120 μg ml^−1^ spectinomycin or 10 μg ml^−1^ erythromycin, respectively. Plates as well as liquid cultures were incubated at 37 °C and 5 % CO_2_.

### General DNA Techniques

Commercial kits, following the manufacturer’s protocols were used to isolate DNA (GeneEluteTM Bacterial Genomic DNA, Sigma-Aldrich; QIAamp® DNA Mini Kit, Qiagen, Hilden, Germany) or plasmids (QIAprep® Spin Miniprep Kit, Qiagen). Polymerase chain reaction (PCR) was conducted using standard protocols for *Taq* polymerase (Roche, Mannheim, Germany). Default settings for PCR were an initial denaturation at 95 °C for 5 min. Followed by 32 cycles of 1 min at 95 °C, 30 s at 50 °C, 1-3 min at 72 °C rounded up by a final elongation of 7 min at 72 °C. For templates longer than 3000 bp the Expand Long Template PCR system: DNA pol. Mix (Roche) was used. Clean up of PCR products was performed with NucleoSpin® Gel and PCR Clean-up (Machery-Nagel, Düren, Germany). Primers used in this study are listed in supplementary Table S2. Commercial nucleotides sequencing was carried out by Eurofins Genomics (Ebersberg, Germany) and Microsynths Seqlab laboratories (Göttingen, Germany).

### *Sil* and *blp3* Screen

For obtaining the *sil* and *blp3* sequence information, primers were designed for this region based on the genome sequence of *S. anginosus*_1_2_62CV (Accession: KY315455.1). PCRs were performed on the chromosomal DNA of *S. anginosus* strains BSU 1211, BSU 1326, BSU 1370 and BSU 1401 using primer pairs 1/2, 3/4 or 3/7, 5/6, 8/9, 10/17, 11/16, 12/13, 14/15. All PCR products were purified and sequenced. (Accession numbers: BSU1211: MZ766502, BSU1326: MZ766503, BSU1370: MZ766504, BSU1401: MZ766505)

### Radial Diffusion Assay

To measure bacteriocin activity, a modified radial diffusion assay (RDA) was conducted^70^. For testing antagonism of bacteria against other bacteria a one-layer RDA was performed, whereas for investigating the antimicrobial activity of supernatant or peptides a two-layer RDA was done. In both cases, overnight cultures of target strains as well as putative bacteriocin producer strains (BPS) were centrifuged at 3000 x g for 10 min and the pellet was solved in 10 mM phosphate buffer. After repeated centrifugation at 3000 x g for 10 min the pellet was reconstituted in 5 ml 10 mM phosphate buffer. Following O.D._600 nm_ determination each plate was seeded with 2 x 10^7^ bacterial cells of the target strains.

For a one-layer RDA bacterial cells were inoculated in still liquid Trypticase Soy Agar and 20 ml plates were poured. After solidification, holes were cut into the agar with sterile wide bore pipette tips (Axygen–a corning brand). These wells were filled with overnight cultures of putative BPS, which were adjusted to an O.D._600 nm_ of 0.5. Following overnight incubation at 37 °C and 5 % CO_2_, inhibition zones were measured in cm. In some assays 100 ng of the autoinducing peptide SilCR_SAG-C_ (GWLEDLFSPYFKKYKLGKLGQPDLG) were added simultaneously with the bacteria to test its effect on bacteriocin production. The peptide was synthesized by the Core facility of functional peptidomics (UPEP, Ulm University, Ulm, Germany) based on the deduced sequence of SilCR_SAG-C_ of *S. anginosus* strain BSU 1211. Purity levels of SilCR_SAG-C_ exceeded 95% and were determined via high performance liquid chromatography (HPLC).

To carry out a two-layer RDA bacterial cultures were inoculated in 15 ml of still liquid 1 % agarose and were poured into a petri dish. Wells were punched into the solidified agarose and filled with the test substance. After 3h of aerobic incubation at 37 °C an overlay with 10 ml Trypticase Soy Agar was performed. Following solidification plates were incubated at 37 °C and 5 % CO_2_ overnight. Inhibition zones were quantified in cm. This assay was used to assess the antimicrobial activity of differently treated cell free supernatant (CFS). Furthermore, the peptide sequence of Angicin was deduced from *S. anginosus* BSU 1211 DNA sequence and in a next step the 54 amino acid sequence without leader peptide (GSGYCKPVMVGANGYACRYSNGRWDYKVTKGIFQATTDVIVKGWAEYGPWIPRH) was synthesized by Peptide Specialty Laboratories GmbH (PSL GmbH, Heidelberg, Germany). Angicin was purified using HPLC and afterwards purity was analyzed with HPLC and MS-MALDI. The determined purity was >98%. The molecular mass was calculated at 6053.1 g/mol, thus 6.05 μg/ml equal 1 μM. It was solved in ultra-pure water. The activity of this peptide was investigated via a two-layer RDA.

### Construction of mutants

### Construction of *blp3.3* mutants

Based on the plasmid pAT28 two different *blp3.3* constructs were created and cloned into *E. coli* EC101. One vector contained *blp3.3* alone and a second vector encoded *blp3.3* with its endogenous promoter (*promblp3.3*). *Blp3.3* without promoter was amplified with primers 18/20 and *promblp3.3* with primers 19/20. All primers that were used introduced EcoRI (Roche) and BamHI (Roche) restriction sites, for cloning purposes. In a next step these constructs were transformed into *S. anginosus* strain SK52 and *S. constellatus* BSU 1213 by electroporation as described elsewhere ^71^. In brief, bacterial cells were made competent by several washing steps in 10 % glycerol. Then 1 μg of plasmid DNA was added and cells were exposed to an electrical pulse. Clones were selected on THY-plates containing 120 μg ml^−1^ spectinomycin. Correct insertions were confirmed by nucleotide sequencing.

### Construction of *blp3* and Angicin deletion mutants

A *S. anginosus* BSU 1211Δ*blp3* strain was created via splicing by overlap extension PCR (SOE) and competence related transformation as described in Bauer et al.^72^. In brief, *blp3* flanking regions of *S. anginosus* BSU 1211 were amplified with primer pairs 21/22 for Thereby, an overlap t1 (F1) and 23/24 for Fragment2 (F2). The primers additionally introduced an overlap to either a lox66 or a lox71 site. The spectinomycin resistance gene was amplified from pGA14-Spc with primers 25/26, which also introduced a lox66 or lox71 site, respectively, adjacent to the spectinomycin gene. Thereby, an overlap of all fragments was achieved. All fragments were fused via SOE-PCR. Subsequently, the linear DNA construct was transformed into *S. anginosus* strain BSU 1211 by inducing natural competence with CSP-1^72^. BSU 1211 was incubated with 100 ng CSP-1 and after 40 min the DNA construct was added. After two hours of incubation bacteria were plated on THY-spectinomycin plates. To eliminate the spectinomycin resistance gene, spectinomycin positive clones were transformed with a cre-recombinase containing plasmid (pAT18-cre-rectufA), which recombines the two lox-sites to a singular lox72 site. By incubating the resulting clones without antibiotics, plasmid loss was induced to create a markerless deletion strain.

The same method was used for creating single gene deletion mutants of the *blp3* region. For a deletion of *blp3.1* primers 5/27 (F1) and 28/29 (F2), for *blp3.4* primers 7/30 (F1) and 31/3 (F2), for *blp.3.6* primers 32/33 (F1) and 34/35 (F2) were used.

To control a successful transformation all constructs were sequenced therefore primers 1/5 (*blp3*) 36/37 (*blp3.1*), 38/39 (*blp3.4*) and 40/41 (*blp3.6*) were applied.

### Complementation of *S. anginosus* BSU 1211Δ*blp3*

To complement the deletion mutant, *S. anginosus* BSU 1211Δ*blp3* was transformed with pAT18-*blp3* by using the natural competence system as previously described. The *blp3* region was amplified using primers 42/43 and cloned into pAT18. Transformed cells were plated on sheep blood agar plates containing 10 μg ml^−1^ erythromycin and incubated anaerobically at 37 °C.

### Cell free supernatant (CFS)

Putative bacteriocin producing strains were grown in THY supplemented with 10 % FBS for 24 h. After centrifugation for 30 min at 4 °C and either 4000 x g (50 ml) or 11 900 x g (UZ) (350 ml) a sterile filtration was performed. Cell free supernatant (CFS) was precipitated with 35 % Ammonium sulfate under stirring conditions for 1 h at 4 °C. Following centrifugation for 30 min, at 4 °C and 4000 x g the supernatant was discarded, and the pellet was resuspended in 10 mM phosphate buffer in one tenth of the original culture volume. In a next step, samples were concentrated via 3K filters (Amicon^®^ Ultra Centrifugal Filters). The concentrate contained the active fraction of the peptide. CFS activity was afterwards tested via a two-layer radial diffusion assay.

### Effect of environmental factors on Angicin activity

The stability of bacteriocins was investigated in regard to pH, temperature, degradation by enzymes and time. CFS was adjusted to pH values ranging from 2 to10. Following overnight incubation at the adjusted pH at room temperature (RT), pH was neutralized and the CFS tested for antibacterial activity. Furthermore, CFS was incubated at 40, 60, 80, 90 °C for 1 h or at 90 and 100 °C for 30 min. After cooling down to RT activity was measured. To assess bacteriocin degradation either proteinase K or Lipase of *Aspergillus niger* was added to CFS to a final concentration of one mg/ml and incubated for 1 h at 37 °C. After inactivation of the enzymes for 1 h at 70 °C and cooling to RT antimicrobial activity was quantified. To further assess stability at low temperatures CFS was stored at 4 °C, −20 °C and −70 °C for one year and then inhibitory activity was analyzed. Activity of 50 μg/ml Angicin was tested after incubation at 60, 80 and 90 °C for 1 h. Furthermore, treatment of 5 μg/ml Angicin with 1 mg/ml lipase or proteinase K was conducted and afterwards activity was tested. All measurements were carried out in triplicate. If the test compound was inactive and not used as a control only two independent experiments were conducted.

### Inhibition assay

In a 96-well plate a serial dilution of CFS from either *S. anginosus* BSU 1211 or the deletion mutant was prepared in THY broth ranging from 100 % CFS to 6.25 %. THY broth without CFS supplementation served as control. 5 μl of target bacteria, adjusted to an O.D._600 nm_ of 0.1, were added to each well in a final volume of 100 μl. The plate was incubated at 37 °C and 5 % CO_2_ for 24 h. Absorbance _600 nm_ was measured every hour for 9 h and after 24 h in a Tecan microplate reader M infinite 200. At least five independent experiments were conducted. Measurements were done in triplicates.

### Survival assay

An overnight culture of *L. monocytogenes, L. ivanovii* or *L. grayi* was diluted to an O.D._600 nm_ of 0.02 in 10 ml THY. When an O.D._600 nm_ of 0.1 was reached, 1 ml was transferred to a 1.5 ml Eppendorf tube and centrifuged at 8800 x g for 2 min. The supernatant was discarded and the pellet reconstituted in 750 μl 10 mM phosphate buffer with 10 % Tryptic Soy broth (TSB). 250 μl CFS of either *S. anginosus* BSU 1211 or *S. anginosus* BSU 1211Δ*blp3* was added and mixed. Cells were incubated at 37 °C and after 0 h, 30 min, 1 h and 2 h serial dilutions were performed and plated on THY. After overnight incubation at 37 °C and 5 % CO_2_ colony forming units per ml were determined. At least five independent experiments were performed with technical duplicates.

### Electron microscopy

Transmission electron microscopy was performed to investigate the effect of active CFS on target bacteria. Therefore, overnight cultures of target bacteria (*S. constellatus, L. monocytogenes, L. ivanovii* and *L. grayi*) were adjusted to an O.D._600nm_ of 0.02 and grown to an O.D._600nm_ of 0.1 and then 1 ml was transferred into a 1.5 ml Eppendorf tube and centrifuged at 8800 x g for 2 min. The pellet was resuspended in 750 μl 10 mM phosphate buffer with 10 % TSB. 25 % (250 μl) CFS from either the *S. anginosus* BSU 1211 or *S. anginosus* BSU 1211Δ*blp3* were added and subsequently incubated at 37 °C. *S. constellatus* and *L. monocytogenes* were incubated with CFS for 2 h. In contrast, *L. grayi* and *L. ivanovii* were only incubated for 15 min, since killing of target cells happend very fast. After centrifugation at 8800 x g for 2 min the supernatant was removed. Cells were reconstituted in 40 μl 10 mM phosphate buffer with 10 % TSB. Then 40 μl of double concentrated fixing solution was added, consisting of 3.5% glutharaldehyde, 1% Saccharose in phosphate buffer). Afterwards samples were postfixed in osmium tetroxide, dehydrated in a graded series of propanol, embedded in Epon, and ultrathin sectioned according to standard procedures. Samples were analyzed with a Jeol 1400 Transmission Electron Microscope. At least 20 images per sample were taken. Experiment was conducted once.

### Recombinant expression of the immunity protein Blp3.3

Blp3.3 protein was recombinantly expressed in *Escherichia coli* RV308. The coding region of Blp3.3 was synthesized (Eurofins) and cloned to the C terminus of a 11 His-tagged maltose-binding protein in a pMAL-C5X vector (New England Biolabs) separated by a cleavage site for tobacco etch virus protease. Protein expression was performed in mineral salt medium M9 by addition of 1 mM IPTG. Protein purification was done in eight steps: (1) amylose resin high flow (New England Biolabs) chromatography using a step elution from 0 mM to 10 mM maltose in Tris buffer A (20 mM Tris/HCl, pH 7.5, 200 ml NaCl), (2) Ni Sepharose fast flow resin (GE Healthcare) chromatography using a step elution from 50 mM to 200 mM Imidazol in Tris buffer B (20 mM Tris/HCl, pH 8.0, 150 mM NaCl), (3) fusion protein cleavage by overnight incubation with tobacco etch virus protease at 34 °C, (4) Ni Sepharose fast flow resin chromatography to separate Blp3.3 from the fusion protein and maltose-binding protein, (5) Source 15 RPC resin (GE Healthcare) chromatography using a linear gradient from 0 % to 86 % (v/v) of acetonitrile in 0.1 % (v/v) trifluoroacetic acid, (6) lyophilization with an alpha 2-4 LD plus freeze dryer (Christ) and redissolving in Tris buffer B, (7) Superdex 75 resin (GE Healthcare) chromatography using an isocratic elution in Tris buffer B and (8) Source 15 RPC resin chromatography (same condition as 5). The purified Blp3.3 was lyophilized and stored at −80 °C until further use. The chemical identity of Blp3.3 was verified by electrospray ionisation mass spectrometry. The determined monoisotopic mass (11247.0 Da) corresponds well to the theoretical monoisotopic mass expected from the amino acid sequence (11246.9 Da).

### SYTOX Green Membrane Permeabilization Assay

A SYTOX Green Membrane Permeabilization Assay was used to assess the effect of synthesized Angicin against *L. monocytogenes.* Therefore, an overnight culture was adjusted to an O.D._600 nm_ of 0.05 and incubated at 37 °C and 5 % CO_2_ till an O.D. 600 nm of 0.1 was reached. One ml of the bacterial culture was transferred into an Eppendorf tube and centrifugation at 10.000 x g for two minutes followed. The pellet was solved in one volume of 10 mM phosphate solution with 0.2 μM Sytox. 90 μl of the bacteria-Sytox solution were mixed with Angicin with final concentrations ranging from 100 μg/ml to 6.25 μg/ml. A Tecan microplate reader M infinite 200 reader was used to measure fluorescence intensity. Excitation/Emission wavelength were 488/530 nm, respectively. Cells treated with 70 % Ethanol for five minutes were used as positive control. Measurements were performed in triplicates and in five independent measurements.

### Bioinformatic and statistical analysis

As a source for nucleotide sequences the GenBank database (http://www.ncbi.nlm.nih.gov/) was used. Homology searches were conducted using Basic Local Alignment Search Tool (http://www.ncbi.nlm.nih.gov/Blast/)^23^. For further genetic investigations SnapGene 5.0 (https://www.snapgene.com/) was accessed. Genetic alignments were done using CLC Main Workbench v7.7.3 (http://www.clcbio.com) with default settings (gap open cost value 10.0, gap extension cost value 1.0). Bioinformatical analysis of the *blp3* region was done with Bagel4 (http://bagel4.molgenrug.nl/)^73^. Analysis of Angicin peptide sequence was performed with Bachem (https://www.bachem.com/service-support/peptide-calculator/). Endogenous promotor search was carried out with Softberry-BPROM (http://www.softberry.com/berry.phtml?topic=bprom&group=programs&subgroup=gfindb). GraphPad Prism V6 was used to create graphs and do statistical analysis. If not otherwise specified, all experiments were conducted independently at least five times. Experiments with inactive compounds were repeated at least twice. Depicted is always mean ± standard deviation.

## Supporting Information

Figure S1: Genetic alignment of amino acid sequences of Angicin prepetide and homologs, Figure S2: Activity of cell free supernatant, Figure S2: Growth inhibition of *Listeria monocytogenes* by CFS treatment, Figure S3: Effect of recombinantly expressed Blp3.3 on antimicrobial activity, Table S1+2: Bacterial strains, plasmids and primers used in this study.

## Acknowledgments

RB, BS, PW and MF are supported by the German Research Foundation (DFG) within the CRC 1279, subprojects A02 and A03. Work of GMS was supported by the Novo Nordisk Fonden within the framework of the Fermentation-based Biomanufacturing Initiative (FBM) (FBM-grant: NNF17SA0031362).

VV thanks the International Graduate School in Molecular Medicine Ulm for the support. We thank M. Wunderlin (Service center mass spectrometry, Ulm University) for mass spectrometric analysis of Blp3.3. The authors like to express thanks to Prof. Dr. Riedel, RWTH Aachen University, University Hospital Düsseldorf and INSTAND e.V. for providing microorganisms.

## Author Contributions

VV, RB and BS designed this study. VV and SM conducted the experiments. NS and CH expressed Blp3.3 under the supervision of MF. GS helped designing experiments to get active cell free supernatant. PW helped in the preparation and analysis of TEM samples. VV and BS analyzed the results. VV wrote and BS reviewed this manuscript.

## References

1. Poole, P. M. & Wilson, G. Occurrence and cultural features of Streptococcus milleri in various body sites. J. Clin. Pathol. 32, 764–768 (1979).

2. Whiley, R. A., Beighton, D., Winstanley, T. G., Fraser, H. Y. & Hardie, J. M. Streptococcus intermedius, Streptococcus constellatus, and Streptococcus anginosus (the Streptococcus milleri group): association with different body sites and clinical infections. J. Clin. Microbiol. 30, 243–244 (1992).

3. Kobo, O., Nikola, S., Geffen, Y. & Paul, M. The pyogenic potential of the different Streptococcus anginosus group bacterial species: retrospective cohort study. Epidemiol. Infect. 145, 3065–3069 (2017).

4. Laupland, K. B., Ross, T., Church, D. L. & Gregson, D. B. Population-based surveillance of invasive pyogenic streptococcal infection in a large Canadian region. Clin. Microbiol. Infect. Off. Publ. Eur. Soc. Clin. Microbiol. Infect. Dis. 12, 224–230 (2006).

5. Reissmann, S. et al. Contribution of Streptococcus anginosus to infections caused by groups C and G streptococci, southern India. Emerg. Infect. Dis. 16, 656–663 (2010).

6. Sibley, C. D. et al. McKay agar enables routine quantification of the ‘Streptococcus milleri’ group in cystic fibrosis patients. J. Med. Microbiol. 59, 534–540 (2010).

7. Siegman-Igra, Y., Azmon, Y. & Schwartz, D. Milleri group streptococcus--a stepchild in the viridans family. Eur. J. Clin. Microbiol. Infect. Dis. Off. Publ. Eur. Soc. Clin. Microbiol. 31, 2453–2459 (2012).

8. Kurm, V. et al. Competition and predation as possible causes of bacterial rarity. Environ. Microbiol. 21, 1356–1368 (2019).

9. Alvarez-Sieiro, P., Montalbán-López, M., Mu, D. & Kuipers, O. P. Bacteriocins of lactic acid bacteria: extending the family. Appl. Microbiol. Biotechnol. 100, 2939–2951 (2016).

10. Arnison, P. G. et al. Ribosomally synthesized and post-translationally modified peptide natural products: overview and recommendations for a universal nomenclature. Nat. Prod. Rep. 30, 108–160 (2013).

11. Mendonca, M. L. et al. The sil Locus in Streptococcus Anginosus Group: Interspecies Competition and a Hotspot of Genetic Diversity. Front. Microbiol. 7, 2156 (2016).

12. Miller, E. L. et al. Eavesdropping and crosstalk between secreted quorum sensing peptide signals that regulate bacteriocin production in Streptococcus pneumoniae. ISME J. 12, 2363–2375 (2018).

13. Ni, J. et al. Autoregulation of lantibiotic bovicin HJ50 biosynthesis by the BovK-BovR two-component signal transduction system in Streptococcus bovis HJ50. Appl. Environ. Microbiol. 77, 407–415 (2011).

14. Shanker, E. & Federle, M. J. Quorum Sensing Regulation of Competence and Bacteriocins in Streptococcus pneumoniae and mutans. Genes 8, (2017).

15. Barbour, A., Philip, K. & Muniandy, S. Enhanced production, purification, characterization and mechanism of action of salivaricin 9 lantibiotic produced by Streptococcus salivarius NU10. PloS One 8, e77751 (2013).

16. Hertzog, B. B. et al. A Sub-population of Group A Streptococcus Elicits a Population-wide Production of Bacteriocins to Establish Dominance in the Host. Cell Host Microbe 23, 312–323.e6 (2018).

17. Soto, C., Padilla, C. & Lobos, O. Mutacins and bacteriocins like genes in Streptococcus mutans isolated from participants with high, moderate, and low salivary count. Arch. Oral Biol. 74, 1–4 (2017).

18. Belotserkovsky, I. et al. Functional analysis of the quorum-sensing streptococcal invasion locus (sil). PLoS Pathog. 5, e1000651 (2009).

19. Hidalgo-Grass, C. et al. A locus of group A Streptococcus involved in invasive disease and DNA transfer. Mol. Microbiol. 46, 87–99 (2002).

20. Founou, R. C., Founou, L. L. & Essack, S. Y. Clinical and economic impact of antibiotic resistance in developing countries: A systematic review and meta-analysis. PLOS ONE 12, e0189621 (2017).

21. Mulani, M. S., Kamble, E. E., Kumkar, S. N., Tawre, M. S. & Pardesi, K. R. Emerging Strategies to Combat ESKAPE Pathogens in the Era of Antimicrobial Resistance: A Review. Front. Microbiol. 10, 539 (2019).

22. van Heel, A. J. et al. BAGEL4: a user-friendly web server to thoroughly mine RiPPs and bacteriocins. Nucleic Acids Res. 46, W278–W281 (2018).

23. Altschul, S. F., Gish, W., Miller, W., Myers, E. W. & Lipman, D. J. Basic local alignment search tool. J. Mol. Biol. 215, 403–410 (1990).

24. Nissen-Meyer, J., Oppegård, C., Rogne, P., Haugen, H. S. & Kristiansen, P. E. Structure and Mode-of-Action of the Two-Peptide (Class-IIb) Bacteriocins. Probiotics Antimicrob. Proteins 2, 52–60 (2010).

25. Bogaardt, C., van Tonder, A. J. & Brueggemann, A. B. Genomic analyses of pneumococci reveal a wide diversity of bacteriocins - including pneumocyclicin, a novel circular bacteriocin. BMC Genomics 16, 554 (2015).

26. Hossain, M. S. & Biswas, I. Mutacins from *Streptococcus mutans* UA159 Are Active against Multiple Streptococcal Species. Appl. Environ. Microbiol. 77, 2428–2434 (2011).

27. Ovchinnikov, K. V. et al. Novel Group of Leaderless Multipeptide Bacteriocins from Gram-Positive Bacteria. Appl. Environ. Microbiol. 82, 5216–5224 (2016).

28. Remschmidt, C. et al. Continuous increase of vancomycin resistance in enterococci causing nosocomial infections in Germany ? 10 years of surveillance. Antimicrob. Resist. Infect. Control 7, 54 (2018).

29. Aas, J. A., Paster, B. J., Stokes, L. N., Olsen, I. & Dewhirst, F. E. Defining the normal bacterial flora of the oral cavity. J. Clin. Microbiol. 43, 5721–5732 (2005).

30. Paster, B. J., Olsen, I., Aas, J. A. & Dewhirst, F. E. The breadth of bacterial diversity in the human periodontal pocket and other oral sites. Periodontol. 2000 42, 80–87 (2006).

31. Dawid, S., Roche, A. M. & Weiser, J. N. The blp bacteriocins of Streptococcus pneumoniae mediate intraspecies competition both in vitro and in vivo. Infect. Immun. 75, 443–451 (2007).

32. Kommineni, S. et al. Bacteriocin production augments niche competition by enterococci in the mammalian gastrointestinal tract. Nature 526, 719–722 (2015).

33. Drucker, D. B. & McKillop, C. M. Bacteriocin production by Streptococcus milleri. Can. J. Microbiol. 28, 278–283 (1982).

34. Willcox, M. D. & Drucker, D. B. Partial characterisation of the inhibitory substances produced by Streptococcus oralis and related species. Microbios 55, 135–145 (1988).

35. García-Curiel, L., del Rocío López-Cuellar, Ma., Rodríguez-Hernández, A. I. & Chavarría-Hernández, N. Toward understanding the signals of bacteriocin production by Streptococcus spp. and their importance in current applications. World J. Microbiol. Biotechnol. 37, 15 (2021).

36. Lozo, J. et al. Molecular Characterization of a Novel Bacteriocin and an Unusually Large Aggregation Factor of Lactobacillus paracasei subsp. paracasei BGSJ2-8, a Natural Isolate from Homemade Cheese. Curr. Microbiol. 55, 266–271 (2007).

37. Tosukhowong, A. et al. Garvieacin Q, a novel class II bacteriocin from Lactococcus garvieae BCC 43578. Appl. Environ. Microbiol. 78, 1619–1623 (2012).

38. Tymoszewska, A., Walczak, P. & Aleksandrzak-Piekarczyk, T. BacSJ—Another Bacteriocin with Distinct Spectrum of Activity that Targets Man-PTS. Int. J. Mol. Sci. 21, 7860 (2020).

39. Whitford, M. F., McPherson, M. A., Forster, R. J. & Teather, R. M. Identification of bacteriocin-like inhibitors from rumen Streptococcus spp. and isolation and characterization of bovicin 255. Appl. Environ. Microbiol. 67, 569–574 (2001).

40. Tymoszewska, A., Diep, D. B., Wirtek, P. & Aleksandrzak-Piekarczyk, T. The Non-Lantibiotic Bacteriocin Garvicin Q Targets Man-PTS in a Broad Spectrum of Sensitive Bacterial Genera. Sci. Rep. 7, 8359 (2017).

41. Cookson, A. L., Noel, S. J., Kelly, W. J. & Attwood, G. T. The use of PCR for the identification and characterisation of bacteriocin genes from bacterial strains isolated from rumen or caecal contents of cattle and sheep. FEMS Microbiol. Ecol. 48, 199–207 (2004).

42. Cao-Hoang, L., Marechal, P. A., Le-Thanh, M. & Gervais, P. Synergistic action of rapid chilling and nisin on the inactivation of Escherichia coli. Appl. Microbiol. Biotechnol. 79, 105–109 (2008).

43. Heesterbeek, D. A. C. et al. Complement-dependent outer membrane perturbation sensitizes Gram-negative bacteria to Gram-positive specific antibiotics. Sci. Rep. 9, 3074 (2019).

44. Li, Q., Montalban-Lopez, M. & Kuipers, O. P. Increasing the Antimicrobial Activity of Nisin-Based Lantibiotics against Gram-Negative Pathogens. Appl. Environ. Microbiol. 84, (2018).

45. Stevens, K. A., Sheldon, B. W., Klapes, N. A. & Klaenhammer, T. R. Nisin treatment for inactivation of Salmonella species and other gram-negative bacteria. Appl. Environ. Microbiol. 57, 3613–3615 (1991).

46. Vasilchenko, A. S. & Valyshev, A. V. Pore-forming bacteriocins: structural–functional relationships. Arch. Microbiol. 201, 147–154 (2019).

47. Chakchouk-Mtibaa, A. et al. Safety Aspect of *Enterococcus faecium* FL31 Strain and Antibacterial Mechanism of Its Hydroxylated Bacteriocin BacFL31 against *Listeria monocytogenes*. BioMed Res. Int. 2018, 1–10 (2018).

48. Wayah, S. B. & Philip, K. Characterization, yield optimization, scale up and biopreservative potential of fermencin SA715, a novel bacteriocin from Lactobacillus fermentum GA715 of goat milk origin. Microb. Cell Factories 17, (2018).

49. Chikindas, M. L., Weeks, R., Drider, D., Chistyakov, V. A. & Dicks, L. M. Functions and emerging applications of bacteriocins. Curr. Opin. Biotechnol. 49, 23–28 (2018).

50. Algburi, A. et al. Subtilosin Prevents Biofilm Formation by Inhibiting Bacterial Quorum Sensing. Probiotics Antimicrob. Proteins 9, 81–90 (2017).

51. Maldonado-Barragï¿½n, A. et al. Garvicin A, a Novel Class IId Bacteriocin from Lactococcus garvieae That Inhibits Septum Formation in L. garvieae Strains. Appl. Environ. Microbiol. 79, 4336–4346 (2013).

52. Fimland, G., Eijsink, V. G. H. & Nissen-Meyer, J. Comparative studies of immunity proteins of pediocin-like bacteriocins. Microbiology 148, 3661–3670 (2002).

53. Alkhatib, Z., Abts, A., Mavaro, A., Schmitt, L. & Smits, S. H. J. Lantibiotics: how do producers become self-protected? J. Biotechnol. 159, 145–154 (2012).

54. Bastos, M. do C. de F., Coelho, M. L. V. & Santos, O. C. da S. Resistance to bacteriocins produced by Gram-positive bacteria. Microbiol. Read. Engl. 161, 683–700 (2015).

55. Johnsen, L., Fimland, G., Mantzilas, D. & Nissen-Meyer, J. Structure-function analysis of immunity proteins of pediocin-like bacteriocins: C-terminal parts of immunity proteins are involved in specific recognition of cognate bacteriocins. Appl. Environ. Microbiol. 70, 2647–2652 (2004).

56. Nawrocki, K. L., Crispell, E. K. & McBride, S. M. Antimicrobial Peptide Resistance Mechanisms of Gram-Positive Bacteria. Antibiot. Basel Switz. 3, 461–492 (2014).

57. Gursky, L. J. et al. Production of piscicolin 126 by Carnobacterium maltaromaticum UAL26 is controlled by temperature and induction peptide concentration. Arch. Microbiol. 186, 317–325 (2006).

58. Jack, R. W. et al. Characterization of the chemical and antimicrobial properties of piscicolin 126, a bacteriocin produced by Carnobacterium piscicola JG126. Appl. Environ. Microbiol. 62, 2897–2903 (1996).

59. Diep, D. B., Skaugen, M., Salehian, Z., Holo, H. & Nes, I. F. Common mechanisms of target cell recognition and immunity for class II bacteriocins. Proc. Natl. Acad. Sci. U. S. A. 104, 2384–2389 (2007).

60. Martin-Visscher, L. A., Sprules, T., Gursky, L. J. & Vederas, J. C. Nuclear magnetic resonance solution structure of PisI, a group B immunity protein that provides protection against the type IIa bacteriocin piscicolin 126, PisA. Biochemistry 47, 6427–6436 (2008).

61. Barraza, D. E. et al. New insights into enterocin CRL35: mechanism of action and immunity revealed by heterologous expression in Escherichia coli. Mol. Microbiol. 105, 922–933 (2017).

62. Ríos Colombo, N. S., Chalón, M. C., Navarro, S. A. & Bellomio, A. Pediocin-like bacteriocins: new perspectives on mechanism of action and immunity. Curr. Genet. 64, 345–351 (2018).

63. Negash, A. W. & Tsehai, B. A. Current Applications of Bacteriocin. Int. J. Microbiol. 2020, 1–7 (2020).

64. Soltani, S. et al. Bacteriocins as a new generation of antimicrobials: toxicity aspects and regulations. FEMS Microbiol. Rev. 45, fuaa039 (2021).

65. Allerberger, F. & Wagner, M. Listeriosis: a resurgent foodborne infection. Clin. Microbiol. Infect. 16, 16–23 (2010).

66. Perez, R. H., Zendo, T. & Sonomoto, K. Novel bacteriocins from lactic acid bacteria (LAB): various structures and applications. Microb. Cell Factories 13 Suppl 1, S3 (2014).

67. Raza, T., Ullah, S. R., Mehmood, K. & Andleeb, S. Vancomycin resistant Enterococci: A brief review. JPMA J. Pak. Med. Assoc. 68, 768–772 (2018).

68. Zhang, Y., Du, M., Chang, Y., Chen, L. & Zhang, Q. Incidence, clinical characteristics, and outcomes of nosocomial Enterococcus spp. bloodstream infections in a tertiary-care hospital in Beijing, China: a four-year retrospective study. Antimicrob. Resist. Infect. Control 6, 73 (2017).

69. Khan, H. A., Ahmad, A. & Mehboob, R. Nosocomial infections and their control strategies. Asian Pac. J. Trop. Biomed. 5, 509–514 (2015).

70. Maricic, N. & Dawid, S. Using the Overlay Assay to Qualitatively Measure Bacterial Production of and Sensitivity to Pneumococcal Bacteriocins. J. Vis. Exp. (2014) doi:10.3791/51876.

71. Ricci, M. L., Manganelli, R., Berneri, C., Orefici, G. & Pozzi, G. Electrotransformation of Streptococcus agalactiae with plasmid DNA. FEMS Microbiol. Lett. 119, 47–52 (1994).

72. Bauer, R., Mauerer, S., Grempels, A. & Spellerberg, B. The competence system of Streptococcus anginosus and its use for genetic engineering. Mol. Oral Microbiol. (2017) doi:10.1111/omi.12213.

73. van Heel, A. J. et al. BAGEL4: a user-friendly web server to thoroughly mine RiPPs and bacteriocins. Nucleic Acids Res. 46, W278–W281 (2018).

