## Supplemental information for "Angicin, a novel bacteriocin of *Streptococcus anginosus*"

Content

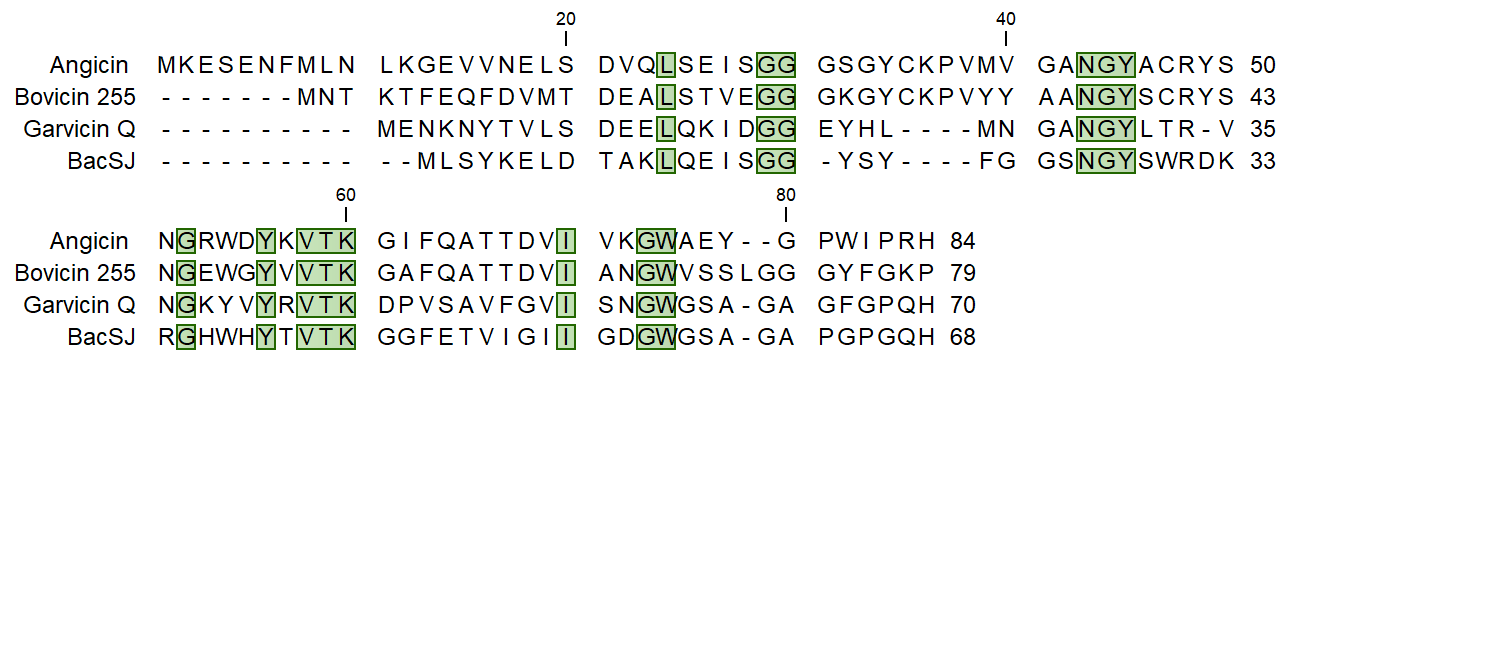

### **Figure S1: Genetic alignment of amino acid sequences of Angicin prepetide and homologs**.

Angicin prepetide was compared to Bovicin variant 255 (GenBank: AAG29818.1), Garvicin Q (GenBank: AEN79392.1) and BacSJ (GenBank: CAR92206.2) using CLC main workbench V7. Conserved residues are marked in green.

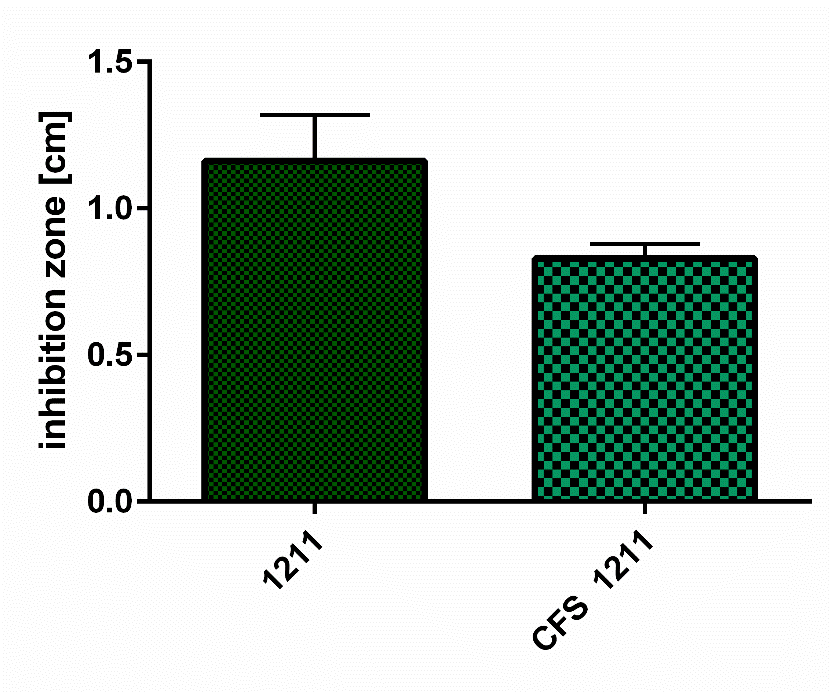

**Figure S2: Activity of cell free supernatant.**

Activity of cell free supernatant (CFS) was assessed in a two-layer RDA against *Listeria monocytogenes* and compared to the activity of *Streptococcus anginosus* BSU 1211 in a one-layer radial diffusion assay.

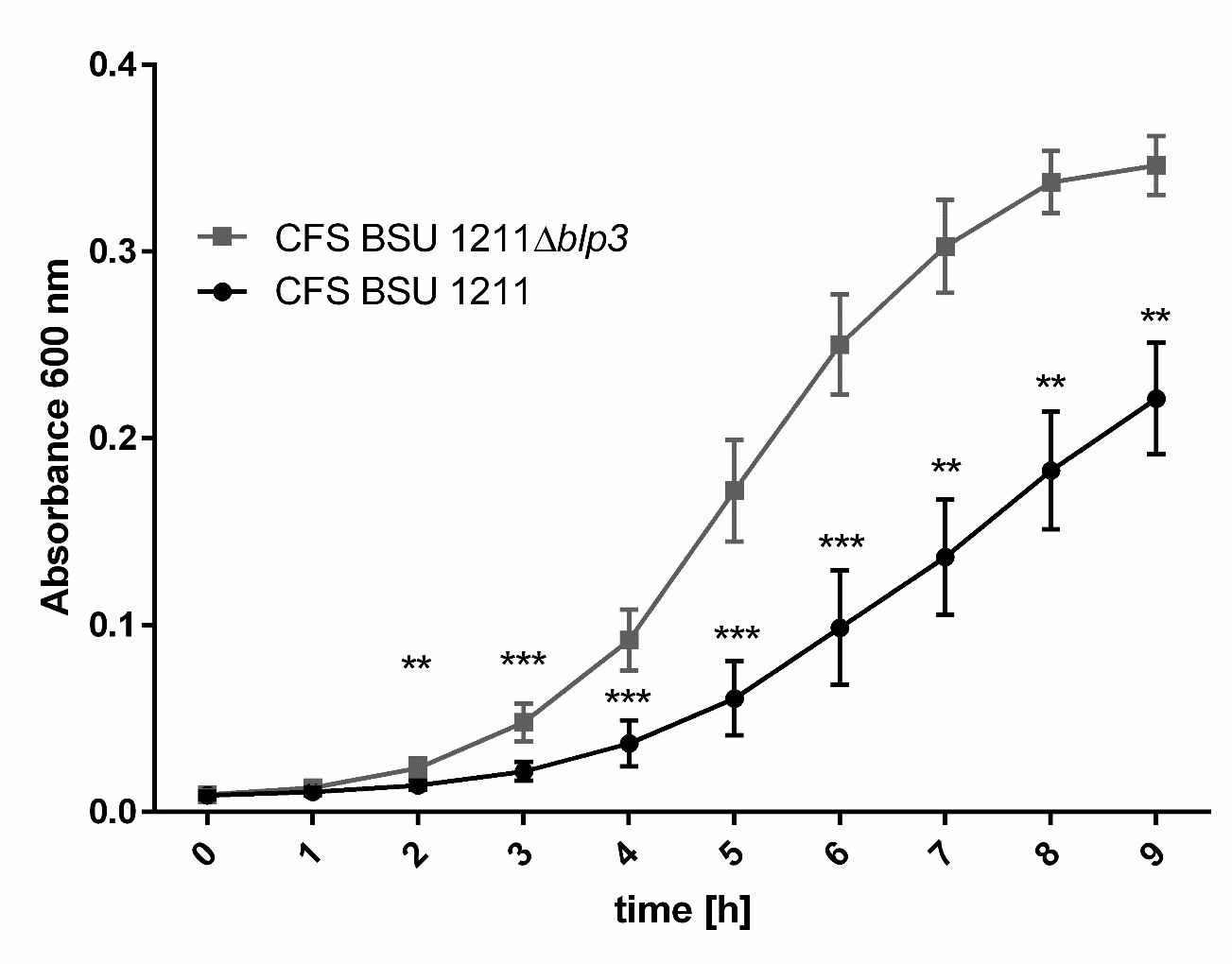

**Figure S3: Growth inhibition of *Listeria monocytogenes* by CFS treatment.**

*L. monocytogenes* was incubated with either cell free supernatant (CFS) of BSU 1211, CFS of BSU 1211∆*blp3* or no CFS over a time course of 9 h. Each hour growth was measured via absorbance at 600 nm in a Tecan plate reader. Data was collected in at least five independent experiments. A significant difference between cells treated with CFS of BSU 1211 or cells treated with CFS of BSU 1211∆*blp3* was tested with a Mann-Whitney-U-test (* indicates p<0.05, ** indicates p<0.01 and *** indicates p<0.001).

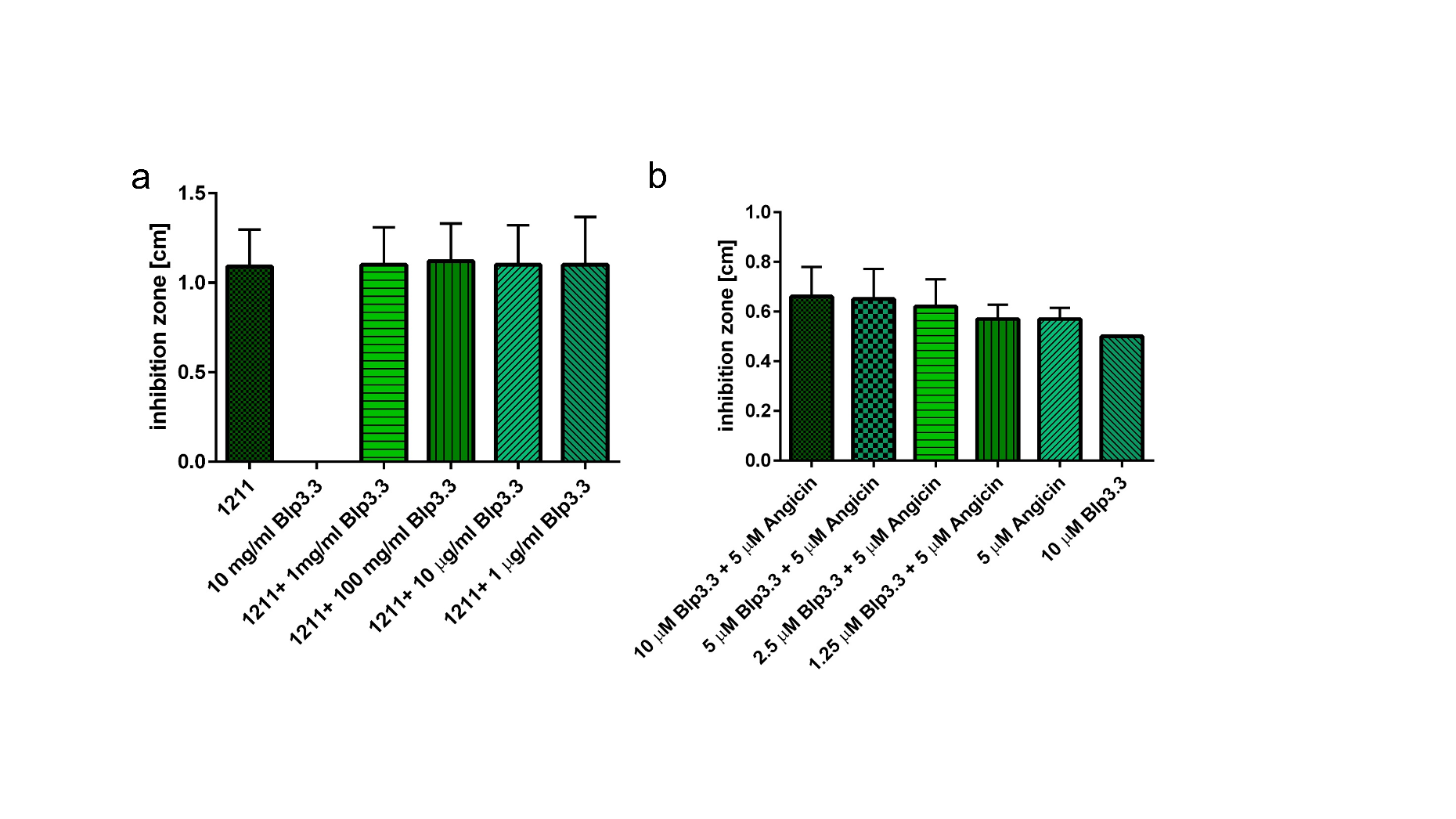

**Figure S4: Effect of recombinantly expressed Blp3.3 on antimicrobial activity.**

(A) Different Blp3.3 concentrations were simultaneously added with BSU 1211 in a one-layer RDA against *L. monocytogenes.* After overnight incubation inhibition zone size was analyzed. (B) Angicin was preincubated with indicated Blp3.3 concentrations for 1 h at 37 °C. Afterwards the antimicrobial activity against *L. monocytogenes* was surveyed in a two-layer RDA. Depicted are five independent experiments conducted with three technical replicates. With a Mann-Whitney-U-test it was controlled for significant differences to either the wildtype or to Angicin alone.

Table S1: Bacterial strains and plasmids used in this study

| **Strain or plasmid** | **Definition** | **Source** |
| --- | --- | --- |
| *Escherichia coli* DH5α | *endA1 hsdR17 supE44* DlacU169(f80lacZDM15) *recA1 gyrA96 thi-1 relA1* | Boehringer |
| *E. coli* EC101 | E. coli JM101 derivative with *repA* from pWV01 integrated into the chromosome | ^1^ |
| *Streptococcus anginosus* BSU 1211^a^ | *S. anginosus*, clinical isolate | ^2^ |
| *S. anginosus* BSU 1324^a^ | *S. anginosus*, clinical isolate | ^2^ |
| *S. anginosus* BSU 1370^a^ | *S. anginosus*, clinical isolate | This study |
| *S. anginosus* BSU 1401^a^ | *S. anginosus*, clinical isolate | ^2^ |
| *S. anginosus* SK 52 | *S. anginosus* type strain, ATCC 33397, Hly+ | ATCC |
| *Streptococcus constellatus* BSU 1213^a^ | *S. constellatus*, clinical isolate | This study |
| *Streptococcus intermedius* BSU 1340^a^ | *S. intermedius*, clinical isolate | This study |
| *Streptococcus pyogenes* BSU 998^a^ | *S. pyogenes* type strain, ATCC 12344 | ATCC |
| *Listeria monocytogenes* EGDe^b^ (BSU 1423) | Ln II Serotype I/2a | ^3^ |
| *Listeria ivanovii* CIP 78.42T^b^ (BSU 1430) | - | ^4^ |
| *Listeria grayi* CIP 68.18T^b^ (BSU 1431) | - | ^4^ |
| *Streptococcus dysagalactiae subsp. equisimilis* BSU 226^c^ | Serotype C | This study |
| *Streptococcus dysagalactiae subsp. equisimilis* BSU 267^c^ | Serotype G | This study |
| *Streptococcus mutans* BSU 269 | DSM 20523 | DSM |
| *Streptococcus agalactiae* BSU 308 | ATCC 12403= NEM 316 | ATCC |
| *Streptococcus suis* BSU 320^d^ | Serotype 2 | This study |
| *Streptococcus porcinus* BSU 852^e^ | - | This study |
| *Streptococcus pneumoniae* BSU 994 | ATCC 49619 | ATCC |
| *Streptococcus mitis* BSU 999 | ATCC 49456 | ATCC |
| *Streptococcus oralis* BSU 1342^a^ | *S. oralis* clinical isolate | This study |
| *Bacillus subtilis* BSU 851 | ATCC 6633 | ATCC |
| *Pseudomonas aeroginosa* BSU 856 | ATCC 27853 | ATCC |
| *Staphylococcus aures* BSU 1348 | MRSA, ATCC 43300 | ATCC |
| *Klebsiella pneumoniae* BSU 1353 | ESBL, ATCC 7000603 | ATCC |
| *Enterococcus faecium* BSU 1516 | VRE, DSM 17050 | DSM |
| *Acinetobacter baumanni* BSU 1514 | *A. baumannii* type strain, ATCC 19606 | ATCC |
| *S. aureus* BSU 995 | ATCC 25923 | ATCC |
| *Staphylococcus epidermidis* BSU 993 | ATCC 12228 | ATCC |
| *S. aureus* BSU 857 | ATCC 29213 | ATCC |
| *S. aureus* BSU 878 | ATCC 13565 | ATCC |
| *Lactobacillus acidophilus* BSU 1314 | *L. acidophilus* | food isolate |
| *Lactobacillus gasseri* BSU 1315 | *L. gasseri* | food isolate |
| *Lactobacillus casei* BSU 1316 | *L. casei* | food isolate |
| *Lactobacillus rhamnosus* BSU 853^e^ | - | This study |
| *Lactobacillus paracasei* BSU 854^e^ | - | This study |
| **Mutants** |  |  |
| *S. anginosus* BSU 1211∆*blp3* |  | This study |
| *S. anginosus* BSU 1211∆*blp3* + pAT18_*blp3* |  | This study |
| *S. anginosus* BSU 1211∆*blp3* + pAT18 |  | This study |
| *S. anginosus* SK52 + pAT18_*blp3* |  | This study |
| *S. anginosus* BSU 1211∆*blp3.1* |  | This study |
| *S. anginosus* BSU 1211∆*blp3.4* |  | This study |
| *S. anginosus* BSU 1211∆*blp3.6* |  | This study |
| *S. anginosus* SK52 + pAT28_*blp3.3* |  | This study |
| *S. anginosus* SK52 + pAT28_*promblp3.3* |  | This study |
| *S. constellatus* BSU 1213 + pAT28_*blp3.3* |  | This study |
| *S. constellatus* BSU 1213 + pAT28_*promblp3.3* |  | This study |
| **Plasmids** |  |  |
| pAT18 | pAT18-lacZα, ori pUC, ori pAmβ1, Em^R^ | ^5^ |
| pAT18-*blp3* | pAT18 derivate carrying the complete *blp3* region, Em^R^ | This study |
| pAT18-cre-rec_tufA_ | pAT18 derivative carrying Cre-recombinase gene under the control of *tufA* promoter, Em^R^ | ^6^ |
| pAT28 | lacZα, ori pUC, ori pAmβ1, Spc^R^ | ^7^ |
| pAT28-EGFP_cfb_ | pAT28 derivate carrying EGFP gene under the control of *cfb* promotor, Spc^R^ | ^8^ |
| pAT28- *blp3.3* | pAT28 derivate carrying *blp3.3* gene, Spc^R^ | This study |
| pAT28- *blp3.3*_prom_ | pAT28 derivate carrying *blp3.3* gene under the control of an endogenous promotor, Spc^R^ | This study |
| pAT28- *blp3.3*_cfb_ | pAT28 derivate carrying *blp3.3* gene under the control of *cfb* promotor, Spc^R^ | This study |
| pGA14-Spc | Replication functions of pWVO1, Em^R^, Spc^R^ | ^9^ |

^a^ isolated at university hospital Ulm, Ulm, Germany

^b^ kindly provided by Prof. Dr. C. Riedel, Ulm University, Ulm, Germany

^c^ kindly provided by RWTH Aachen University, Aachen, Germany

^d^ kindly provided by University Hospital Düsseldorf, Düsseldorf, Germany

^e^ obtained from INSTAND e.V., Düsseldorf, Germany

Table S2: Primers used in this study

| Primer name | Sequence | Nr. |
| --- | --- | --- |
| Blp3_screen1_fwd | tttcataatgttcccgattg | 1 |
| Blp3_screen1_rev | agtatgttaatcgctctaatc | 2 |
| Blp3_screen2_fwd | gggcacagtattgtacagg | 3 |
| Blp3_screen2_rev | tccctaataactagctctac | 4 |
| Blp3_screen3_rev | gtagctctgatagcctgaatg | 5 |
| Blp3.3_fwd | tggtgggcaggaaagaag | 6 |
| Blp3.3_rev | caccctcccagatgttatcg | 7 |
| Sil_screen1_fwd | catagcctctatctggtatatc | 8 |
| Sil_screen1_rev | cgtaaatgacggtctaaataattgg | 9 |
| Sil_screen2_fwd | caattatggcgacgctgatag | 10 |
| Sil_screen2_rev | gcggcgacttatgacaatag | 11 |
| Sil_screen3_fwd | ctgtttcgggagcgactaatc | 12 |
| Sil_screen3_rev | caaatccgtcttaatggaaatg | 13 |
| Sil_screen4_fwd | caatttcacaagcgcgaataatc | 14 |
| Sil_screen4_rev | gtattctactaaccggctttgc | 15 |
| SilCR_1211_fwd | caactatgacgattgcttatg | 16 |
| SilCR_1211_rev | gcagtccatcaccattatc | 17 |
| Blp3.3_1211_EcoRI_fwd | gggcccgaattcatgggaactttaagatgg | 18 |
| Blp3.3_1211_prom_EcoRI_fwd | gggcgcgaattctgtcgctatagtaatggc | 19 |
| Blp3.3_1211_BamHI_rev | ccgggcggatccctagattattctaatatgag | 20 |
| Blp3_1211_F1_fwd | aacgagctgtcttcctgtc | 21 |
| Blp3_1211_F1_rev | gcatacattatacgaacggtacgggtattgctggtggtg | 22 |
| Blp3_1211_F2_fwd | tataatgtatgctatacgaacggtacgactatgaataccaaac | 23 |
| Blp3_1211_F2_rev | aggattggaagttcacaag | 24 |
| lox71_spec_fwd | taccgttcgtatagcatacattatacgaagttatttaaatggcattggtaccc | 25 |
| lox66_spec_rev | taccgttcgtataatgtatgctatacgaagttatatgcctgcaggtcgattttcg | 26 |
| Blp3.1_del_F1_rev | tataatgtatgctatacgaacggtagagtagttgcaggagctgttg | 27 |
| Blp3.1_del_F2_fwd | gcatacattatacgaacggtacacttgccagcatttcagag | 28 |
| Blp3.1_del_F2_rev | ggctggtctgaatccttctac | 29 |
| Blp3.4_del_F1_rev | tataatgtatgctatacgaacggtagaatatggtccgtggattccaag | 30 |
| Blp3.4_del_F2_fwd | gcatacattatacgaacggtagaacatctgatagctcatttacaac | 31 |
| Blp3.6_del_F1_fwd | ctgatagctcatttacaacttc | 32 |
| Blp3.6_del_F1_rev | tataatgtatgctatacgaacggtagggcacagtattgtacagg | 33 |
| Blp3.6_del_F2_fwd | gcatacattatacgaacggtacacttgccagcatttcagag | 34 |
| Blp3.6_del_F2_rev | gaagaacagcatcaatatctc | 35 |
| Blp3.1_del_screen_fwd | cagttatagctgtgcgttgatg | 36 |
| Blp3.1_del_screen_rev | gaagctgaacaacctgattgg | 37 |
| Blp3.4_del_screen_fwd | ccaatcaggttgttcagcttctc | 38 |
| Blp3.4_del_screen_rev | cagtagcaggtggaacgatag | 39 |
| Blp3.6_del_screen_fwd | ctttcctgcccaccagcaaac | 40 |
| Blp3.6_del_screen_rev | gctgcgatatcacgtttatgg | 41 |
| Blp3_1211_EcoRI_fwd | gggcccgaattccaaacgagctgtcttcctg | 42 |
| Blp3_1211_BamHI_rev | ggccgcggatccctgataacactttatagc | 43 |

References

1.	Law, J. *et al.* A system to generate chromosomal mutations in Lactococcus lactis which allows fast analysis of targeted genes. *J. Bacteriol.* **177**, 7011–7018 (1995).

2.	Bauer, R. *et al.* Heterogeneity of Streptococcus anginosus ß-hemolysis in relation to CRISPR/Cas. *Mol. Oral Microbiol.* **35**, 56–65 (2020).

3.	Bécavin, C. *et al.* Comparison of widely used Listeria monocytogenes strains EGD, 10403S, and EGD-e highlights genomic variations underlying differences in pathogenicity. *mBio* **5**, e00969-00914 (2014).

4.	Zetzmann, M. *et al.* DNase-Sensitive and -Resistant Modes of Biofilm Formation by Listeria monocytogenes. *Front. Microbiol.* **6**, 1428 (2015).

5.	Trieu-Cuot, P., Carlier, C., Poyart-Salmeron, C. & Courvalin, P. Shuttle vectors containing a multiple cloning site and a lacZ alpha gene for conjugal transfer of DNA from Escherichia coli to gram-positive bacteria. *Gene* **102**, 99–104 (1991).

6.	Bauer, R., Mauerer, S., Grempels, A. & Spellerberg, B. The competence system of Streptococcus anginosus and its use for genetic engineering. *Mol. Oral Microbiol.* (2017) doi:10.1111/omi.12213.

7.	Trieu-Cuot, P., Carlier, C., Poyart-Salmeron, C. & Courvalin, P. A pair of mobilizable shuttle vectors conferring resistance to spectinomycin for molecular cloning in Escherichia coli and in gram-positive bacteria. *Nucleic Acids Res.* **18**, 4296 (1990).

8.	Aymanns, S., Mauerer, S., van Zandbergen, G., Wolz, C. & Spellerberg, B. High-level fluorescence labeling of gram-positive pathogens. *PloS One* **6**, e19822 (2011).

9.	Smith, H. E., Wisselink, H. J., Vecht, U., Gielkens, A. L. & Smits, M. A. High-efficiency transformation and gene inactivation in Streptococcus suis type 2. *Microbiol. Read. Engl.* **141 ( Pt 1)**, 181–188 (1995).
